# Substrate selectivity and catalytic activation of the type III CRISPR ancillary nuclease Can2

**DOI:** 10.1101/2023.09.18.558242

**Authors:** Kenny Jungfer, Annina Sigg, Martin Jinek

## Abstract

Type III CRISPR-Cas systems provide adaptive immunity against foreign mobile genetic elements through RNA-guided interference. Sequence-specific recognition of RNA targets by the type III effector complex triggers the generation of cyclic oligoadenylate (cOA) second messengers that activate ancillary effector proteins, thus reinforcing the host immune response. The ancillary nuclease Can2 is activated by cyclic tetra-AMP (cA4); however, the mechanisms underlying cA4-mediated activation and substrate selectivity remain elusive. Here we report crystal structures of *Thermoanaerobacter brockii* Can2 (TbrCan2) in substrate- and product-bound complexes. We show that TbrCan2 is a single strand-selective DNase and RNase that binds substrates via a conserved SxTTS active site motif, and reveal molecular interactions underpinning its sequence preference for CA dinucleotides. Furthermore, we identify a molecular interaction relay linking the cA4 binding site and the nuclease catalytic site to enable divalent metal cation coordination and catalytic activation. These findings provide key insights into the molecular mechanisms of Can2 nucleases in type III CRISPR-Cas immunity and may guide their technological development for nucleic acid detection applications.

## INTRODUCTION

Clustered Regularly Interspaced Short Palindromic Repeats (CRISPR) and CRISPR-associated (Cas) systems provide bacteria and archaea with adaptive immunity against invading mobile genetic elements (MGEs) through CRISPR RNA (crRNA)-guided interference (1, 2). CRISPR-Cas systems are categorized into two main classes (class 1 and class 2) that can be further subdivided into six types: type I, III and IV in class 1, and type II, V and VI in class 2 systems (3). In contrast to class 2 systems that deploy single protein effectors during interference, class 1 systems comprise large, multisubunit complexes composed of several Cas proteins that perform different functions. The modularity of class 1 loci gives rise to unique interference mechanisms, which is particularly prevalent in type III CRISPR-Cas immune responses.

During type III CRISPR-Cas interference, the Csm (type III-A, -D and -E) or Cmr (type III-B and - C) effector complex recognizes and degrades foreign-derived RNAs complementary to their crRNA guides (4–6). Binding of cognate target RNAs further activates the Cas10 (Csm1/Cmr2) subunit of the Csm/Cmr complex, which uses its HD nuclease domain to non-specifically degrade single-stranded DNA (ssDNA) (7, 8). In turn, the palm (or cyclase) domain of activated Cas10 converts ATP to 3’-5’ cyclic oligoadenylate (cOA) nucleotides (9, 10). The produced cOAs of system-specific length, commonly cyclic tri-(cA_3_), tetra-(cA_4_) or hexa-adenylates (cA_6_), act as second messengers to stimulate downstream interference pathways by allosterically activating type III CRISPR-Cas ancillary effector proteins that contain CRISPR-associated Rossman-fold (CARF) or SMODS-associated (SAVED) regulatory domains fused to various effector domains (11–13). A broad spectrum of type III-associated CARF and SAVED domain-encoding proteins have been identified that additionally contain effector domains including nucleases, proteases, and transcription factors (14– 16).

The Csm6/Csx1 family of nucleases constitutes the best studied class of ancillary type III effectors. Csm6-like nucleases form homodimeric complexes that harbor C-terminal higher eukaryotic and prokaryotic nuclease (HEPN) domains that non-specifically cleave RNAs in a metal-independent manner upon cOA binding (17–19). The stimulation of Csm6/Csx1 signaling cascades contributes to the inhibition of phage or plasmid proliferation through the induction of cell dormancy or abortive infection. Some Csm6/Csx1 proteins also have an intrinsic ring nuclease activity, thereby limiting their cOA-dependent activation and self-toxicity (20–22). The immune signal amplification afforded by the target-dependent generation of cOAs and induction of Csm6/Csx1 ribonucleases instigated the development of RNA detection tools for diagnostic applications (23–25). However, initial approaches failed to reach sufficient sensitivity in the absence of a prior amplification step, partly due to the autocatalytic inactivation of Csm6. An alternative RNA detection technology with substantially improved detection sensitivity has recently been developed by implementing a new RNA capture approach in combination with the replacement of Csm6 with the CRISPR-associated nuclease 2 (Can2), which lacks ring nuclease activity (26).

The Can2 family is a widely distributed class of type III CRISPR-Cas ancillary effector nucleases that contain PD-(D/E)xK nuclease domains with similarities to type II restriction endonucleases (REase) (27, 28). Can2 form homodimeric complexes that bind a single cOA molecule in a central cavity formed at the dimer interface of the CARF and nuclease domains. Binding to cA_4_, but not to cA_6_, leads to conformational changes that activate the C-terminal REase domain, which results in the degradation of oligonucleotide substrates, conferring immunity against bacteriophages and invading plasmids through growth arrest (28). It remains unclear as to how the cA_4_-induced conformational changes mediate nuclease activation. Interestingly, different Can2 variants exhibit varying nuclease substrate-specificities (26–28). The Can2-like nuclease from *Treponema succinifaciens* TresuCARD1 specifically degrades single-stranded DNA and RNA, whereas Can2 enzymes from *Thioalkalivibrio sulfidiphilus* (TsuCan2) and *Sulfobacillus thermosulfidooxidans* (SthCan2), as well as an archaeal Can2 from Archaeoglobi archaeon JdFR-42 (AaCan2), degrade single-stranded RNA and nick supercoiled plasmid DNA (26–28). Moreover, Can2 nuclease activity exhibits a certain degree of cleavage preference for specific oligonucleotide sequences. AaCan2 was shown to preferentially degrade substrates at poly-A and poly-T sites, whereas TresuCARD1 cleaved upstream of T-(G/A) dinucleotide sequences. However, the molecular basis for substrate preferences in Can2 nucleases remains unclear (26, 28).

Here we present multiple crystal structures of the cA_4_-dependent type III-D ancillary nuclease Can2 from *Thermoanaerobacter brockii* (TbrCan2) in complexes with DNA substrates and products, revealing intricate molecular mechanisms that underpin substrate recognition. Complementing our structural observations with biochemical assays, we identify molecular determinants that account for the substrate selectivity for CA di-nucleotide motifs. Furthermore, we reveal the catalytic activation mechanism of Can2 nucleases involving an allosteric relay from the cA_4_ binding site to the REase catalytic site to enable divalent metal cation coordination and substrate binding.

## MATERIALS AND METHODS

### Protein expression and purification

The sequence of TbrCan2 was amplified from genomic DNA (*Thermoanaerobacter brockii* subsp. *finnii* Ako-1 DSM 1457) and cloned into a Macrolab 16B-series vector (N-terminal hexahistidine tag; TEV protease cleavage site) by ligation-independent cloning. QuikChange site-directed mutagenesis was performed to introduce point mutations. Can2 and its mutants were expressed in Rosetta2 (DE3) *E. coli* cells grown at 37 °C in LB medium. Protein expression was induced at OD_600_ of 0.6 by addition of 0.5 mM IPTG at 18 °C overnight. Cells were harvested and resuspended in 20 mM HEPES-KOH pH 7.5, 500 mM KCl, 15 mM imidazole supplemented with protease inhibitors and lysed using a high-pressure cell lyser (Maximator HPL6). Cell lysates were clarified by centrifugation at 20,000 x g for 30 min and loaded on a 5 ml Ni-NTA Superflow column (QIAGEN) followed by extensive washing. Tagged Can2 was eluted with 20 mM HEPES-KOH pH 7.5, 250 mM KCl, 200 mM imidazole. The imidazole concentration was adjusted to a concentration of 30 mM and the affinity tag was removed in the presence of His-tagged TEV protease overnight at 4 °C. His-tag-containing proteins were removed by reloading the sample on a Ni-NTA affinity column. The untagged protein was concentrated and further purified by size exclusion chromatography (SEC) on a HiLoad 16/600 Superdex 200 column (Cytiva) in 20 mM HEPES-KOH pH 7.5, 250 mM KCl, 1 mM DTT. Pure protein fractions were concentrated and flash-frozen in liquid nitrogen and stored at −80 °C.

### Crystallization and structure determination

For crystallization, TbrCan2 was diluted to a final protein concentration of 150 μM diluted with buffer X (20 mM HEPES-KOH pH 7.5, 100 mM KCl, 1 mM DTT). The TbrCan2^WT^-cA_4_ complex was prepared by mixing 150 μM protein with 225 μM cA_*4*_ in buffer X and assembled at RT for 10 min. The activated TbrCan2^WT/E341A^ complexes in the presence of DNA substrates were assembled by mixing 150 μM of protein in a 1:1.5:3 molar ratio with cA_4_ and substrate DNA oligonucleotide, followed by incubation at 37 °C for 10 min in buffer X supplemented with 20 mM MnCl_2_. Crystals were grown at 20 °C by sitting-drop vapor diffusion by mixing protein samples in a 1:1 ratio (0.2 μl + 0.2 μl) with a reservoir solution (50 mM Tris pH 8, 17.1% PEG MME 550, 4.3% PEG 4000).

Crystals were cryoprotected with 25% PEG 400 and flash-frozen in liquid nitrogen. Diffraction data was collected at the PXI and PXIII beamlines of the Swiss Light Source (Paul Scherrer Institute, Villigen, Switzerland) and processed using the data-processing program package autoPROC (29). Structures were solved by molecular replacement (MR) with Phaser, as part of the PHENIX package (30). For apo Can2, the *in silico* generated AlphaFold2 model of TbrCan2^WT^ was used as a search model (31). The structures of the active Can2 complexes were solved by MR using the refined monomeric apo Can2 structure. Atomic model adjustment and refinement was conducted iteratively using Coot and PHENIX.refine (Supplementary Table 1) (32, 33). Figures were generated with ChimeraX (34).

### Plasmid nuclease assay

*In vitro* nuclease assays with supercoiled M13mp18 RF I dsDNA and M13mp18 ssDNA were performed with 250 nM Can2 in the presence of 2.5 μM cA_4_ and 2 nM circular DNA in reaction buffer (20 mM HEPES pH7.5, 100 mM KCl,5 mM MnCl_2_). The reaction was incubated at 45 °C and was terminated by addition of 10 mM EDTA at indicated time points (control reactions were stopped after 45 min). 6x DNA loading buffer (10 mM Tris pH 7.6, 60 mM EDTA, 60% glycerol, 0.03% bromophenol blue, 0.03% xylene cyanole FF) was added and analyzed by 1% native agarose gel electrophoresis. Gels were scanned with a ChemiDoc Imager (Bio-Rad) and band intensities were quantified with Image Lab 6.1 (Bio-Rad).

### RNase and DNase activity assays

RNA and DNA cleavage assays with linear substrates were carried out with 50 nM 5’-Atto532-labelled DNA and RNA substrates, 250 nM Can2 and 2.5 μM cA_4_ in reaction buffer at 45 °C. Reactions were stopped by addition of 2x denaturing loading dye (5% glycerol, 25 mM EDTA, 90% formamide, 0.15% OrangeG). The reactions were analyzed by 20% denaturing PAGE (20% acrylamide, 7 M urea, 0.5x TBE buffer) and bands were visualized on a Typhoon FLA 9500 laser scanner (GE Healthcare).

For determining *in vitro* nuclease cleavage kinetics under multiple-turnover conditions, 5 nM Can2 was mixed with 50 nM cA_4_ and increasing concentrations of fluorophore-quencher oligonucleotides containing a 5’-6FAM fluorophore and a 3’-Black Hole Quencher 1 (50, 70, 100, 150, 250, 500, 1000, 2000 nM DNA or RNA) in a reaction buffer supplemented with 0.01% TWEEN-20. The triplicate experiments were monitored using a PHERAstar FSX microplate reader (BMG Labtech) measuring excitation and emission wavelength of 485 nm and 520 nm over 35 min at 45 °C. RNA and DNA probes were purchased from Integrated DNA Technologies (IDT).

### Electrophoretic mobility shift assays

For the detection of protein-RNA/DNA interactions, nucleic acid binding assays were performed with 25 nM 5’-Atto532 oligonucleotides and 50 μM cA_4_ in the presence of different amounts of TbrCan2^E341A^. Reactions were incubated in EMSA buffer (20 mM HEPES pH7.5, 25 mM KCl,1 mM DTT, 5 mM EDTA) for 10 min at 37 °C. Samples were loaded on a native 10% polyacrylamide TBE gel and electrophoresis was carried out at 8 °C. Gels were imaged on a Typhoon FLA 9500 and band intensities were quantified as described before. Assays to determine K_D_ were performed in triplicates.

### Analytical SEC cOA binding assays

The binding of TbrCan2 to different cOAs (cA_3_, cA_4_, cA_6_) was assayed by incubating 50 μM ligand with 100 μM (cA_4_) or 300 μM (cA_3_ and cA_6_) TbrCan2 for 10 min at 45 °C in binding buffer (20 mM HEPES pH7.5, 100 mM KCl,5 mM MnCl_2_, 1 mM DTT). Analytical SEC was performed using an ÄKTA™ pure micro (Cytiva). The sample was separated on a Superdex 200 Increase 5/150 GL column that was pre-equilibrated with binding buffer and absorption was detected at a 260 nm and 280 nm wavelength.

### Sequence analysis and construction of phylogenetic tree

A list of homologous sequences was generated by standard protein BLAST (35) using previously published Can2 sequences as query. Redundant sequences were removed using EMBOSS SkipRedundant (36) and CD-HIT (37). The resulting list of 828 Can2 sequences was used as input for multiple sequence alignment using MAFFT v7.429 (38). From this alignment, a phylogenetic tree was generated via PHYLIP neighbor-joining (39). CRISPR system assignment for the CRISPR loci containing *can2* genes was performed with CRISPRCasFinder (40).

## RESULTS

### Phylogenetic analysis of Can2 family nucleases

The Can2 family of type III CRISPR-Cas ancillary effector nucleases is highly abundant in both archaeal and bacterial species. Recent studies have revealed that Can2-like nucleases exhibit distinctive targeting mechanisms despite their shared domain compositions (26–28). Can2 is typically encoded in canonical type III CRISPR-Cas loci, which comprise genes encoding the interference modules and additional accessory effectors, such as the non-specific RNase Csx1 (**Figure 1A**). Phylogenetic analysis of 828 orthologs identified seven distinctive Can2 clades with low sequence identity between them (**Figure 1B**). Analysis of the CRISPR loci that contain the respective *can2* genes shows that the clades often correspond to specific subtypes of type III CRISPR-Cas systems. Clade 5 primarily comprises Can2 proteins from type III-A systems, such as TresuCARD1, while Can2 proteins belonging to clades 2 and 6 originate from type III-B systems, including SthCan2 and TsuCan2 (**Figure 1B**). Clade 1 constitutes the largest cluster and encompasses orthologs that are encoded within type III-D loci (**Figure 1A, B**). Can2 proteins share a conserved domain architecture comprising an N-terminal CARF domain and a C-terminal PD-(D/E)xK nuclease (REase) domain, connected by an α-helical insertion (INS) domain (**Figure 1C**). In contrast, Can2-like nucleases of clade 4 lack the INS domain and only possess a short linker that connects the CARF and nuclease domains (**Figure 1D**). In all clades, Can2 harbors a conserved NxTGGTK motif within the CARF domain, which in SthCan2 and TresuCARD1 has been shown to mediate cyclic oligoadenylate (cOA) binding (**Figure 1D, E**) (28, 41). A conserved E-ExD-(E/S)CK motif, present in most Can2 orthologs, has been shown to be essential for the nuclease activity of TresuCARD1 (28). However, Can2 nucleases from clade 1 instead contain a distinct E-ExD-SxTTS-K motif, suggesting that type III-D Can2 nucleases may differ mechanistically from previously characterized orthologs (**Supplementary Figure S2e**).

**Figure 1.**
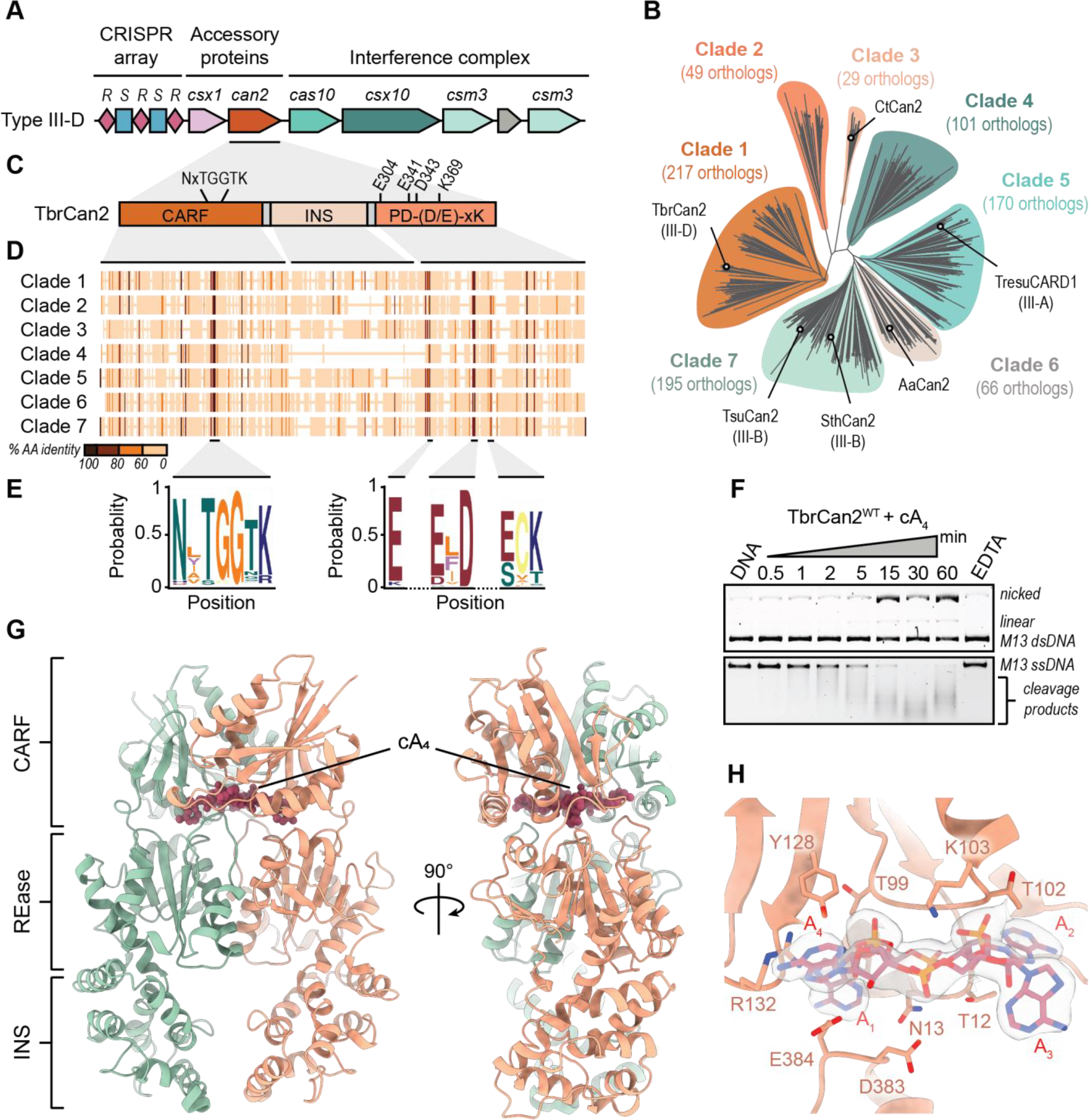
Structure and activity of type III ancillary nuclease TbrCan2. (**A**) Schematic diagram of the *Thermoanaerobacter brockii* finnii Ako-1 type III-D CRISPR-Cas locus; *R*: direct repeat; *S*: spacer. (**B**) Phylogenetic clustering of Can2-like nucleases. (**C**) Schematic diagram of the domain composition of TbrCan2. (**D**) Amino acid sequence conservation among different Can2 clades. (**E**) Sequence logos of conserved motifs within CARF (left) and REase domains (right). (**F**) Nuclease activity assay of TbrCan2 in presence of single-stranded and double-stranded M13 DNA plasmids. (**G**) Crystal structure of TbrCan2 in complex with a cA_4_ ligand at a resolution of 3.2 Å. (**H**) Close-up view of the cA_4_ binding site at the CARF domain interface. 2m*F*_0_-D*F*_C_ composite omit map in **H** is contoured at 1.0 σ and displayed within a radius of 2 Å around the cA_4_ ligand.

### Molecular architecture of the type III-D ancillary nuclease TbrCan2

Previous studies have reported distinct substrate selectivity of Can2 family nucleases (27, 28). Given the distinctive catalytic site composition of clade 1 Can2 orthologs, we chose to characterize the activity and mechanism of the CRISPR type III-D ancillary nuclease Thebr_0943 (TbrCan2; **Figure 1B, C**). TbrCan2 exhibited ssDNase activity *in vitro*, completely degrading single-stranded M13 plasmid DNA in the presence of cA_4_ and MnCl_2_ (**Figure 1F**). Moreover, TbrCan2 nicked supercoiled double-stranded plasmid DNA, albeit with considerably lower efficiency as compared to ssDNA degradation. Nuclease activity was absent in the presence of EDTA, consistent with the divalent metal-dependency of PD-(D/E)xK family nucleases (42).

To gain molecular insights into TbrCan2 activity, we performed X-ray crystallographic studies and initially determined the atomic structure of wild-type TbrCan2 (TbrCan2^WT^) in complex with cA_4_ at a resolution of 3.2 Å (**Figure 1G**). TbrCan2 forms a symmetrical homodimeric complex, binding cA_4_ in a central cavity located between the CARF and REase domains at the dimer interface, similar to other Can2 nucleases (**Supplementary Figure S1**) (27, 28). The CARF domain of TbrCan2 (residues 1-145) exhibits a canonical Rossman fold conserved within the Can2 family, superimposing with TresuCARD1 and SthCan2 CARF domains with root-mean-square deviations (RMSD) of 2.2 Å and 2.3 Å over 139 and 136 residues, respectively (**Supplementary Figure S1b, e**). The cA_4_ molecule is coordinated by an extensive network of hydrogen bonding and hydrophobic interactions involving residues from both the CARF and REase domains. The bases A1 and A3 of cA_4_ are contacted by CARF domain residue Pro16 and REase domain residue Asp383 and Glu384, while nucleobases A2 and A4 are bound by CARF residues Gly11, Thr12, and Ser36 and REase residues Gly410 and Thr409 (**Figure 1H**). The residues Thr99, Gly100, Gly101, Thr102 and Lys103 of the conserved NxTGGTK motif form a network of interactions with the riboses and phosphodiester groups of the cA_4_ ligand, further stabilizing its binding between the two CARF domains. The TbrCan2 REase domain (residues 290-437) displays a similar architecture compared to previously described orthologs, superimposing with a RMSD of 2.3 Å with the REase domains of SthCan2 and TresuCARD1, respectively (**Supplementary Figure S1c, f**). The REase catalytic site in TbrCan2 features conserved residues Glu304, Glu341, Asp343, and Lys369, of which Asp343 directly coordinates a single Mn^2+^ ion in the TbrCan2-cA_4_ complex. In SthCan2 and TresuCARD1, divalent metal binding is additionally mediated by the (E/S)CK REase motif, which is absent in TbrCan2 and replaced with a SxTTS motif that is conserved among clade 1 Can2 orthologs (**Supplementary Figure S1c, f**). Ser356 within this motif, equivalent to Glu308 in CARD1 and Glu291 in SthCan2, is involved in water-mediated, outer-sphere coordination of the Mn^2+^ ion together with Glu304 and Lys369. The catalytic activity of PD-(D/E)xK enzymes occurs via a two-metal ion mechanism (42). Coordination of the second metal ion would be mediated by Asp343 and Glu341 in TbrCan2, based on structural superpositions with type II restriction endonucleases (**Supplementary Figure S9c**). The absence of the second metal ion in the structure implies this binding site has intrinsically low affinity and that ion binding likely occurs cooperatively upon substrate nucleic acid binding.

### TbrCan2 preferentially cleaves single-stranded nucleic acid substrates

To investigate the substrate selectivity of TbrCan2, we analyzed the cOA-dependent nuclease activity of TbrCan2 on linear single-stranded and double-stranded DNA and RNA substrates with similar sequences. A single-stranded DNA (ssDNA) substrate was completely degraded after 30 minutes, while TbrCan2 cleaved the RNA in less than 1 minute, demonstrating potent RNase activity (**Figure 2B**). Double-stranded DNA was cleaved at a low rate, comparable to the background nuclease activity observed in the absence of cA_4_ (**Supplementary Figure S3b, e**), indicating that TbrCan2 primarily targets single-stranded nucleic acids. The observed nuclease activity was dependent on the presence of manganese cations and was significantly reduced in the presence of magnesium (**Figure 2B and Supplementary Figure S3a, d**). Moreover, catalytic activation required the presence of cA_4_, and was not supported by cA_3_ or cA_6_ (**Figure 2B and Supplementary Figure S3b, e**). This was in agreement with analytical size-exclusion chromatography (SEC) binding assays, which revealed that TbrCan2 exhibits stable binding exclusively to cA_4_ (**Supplementary Figure S4**), confirming cA_4_ as the primary activator of TbrCan2. Mutations of conserved residues in the CARF cA_4_ binding site, specifically T12A/N13A and Y128A, did not impede nuclease activation, indicating a certain degree of cooperativity among CARF residues (**Figure 2C**). Alanine substitutions of the catalytic residues Glu304, Asp343, and Lys369 abolished DNase activity. In contrast, RNase activity was partially perturbed by mutations of individual active site residues and was substantially reduced by simultaneous mutations of Glu341 and Lys369 (**Figure 2C and Supplementary Figure S5**).

**Figure 2.**
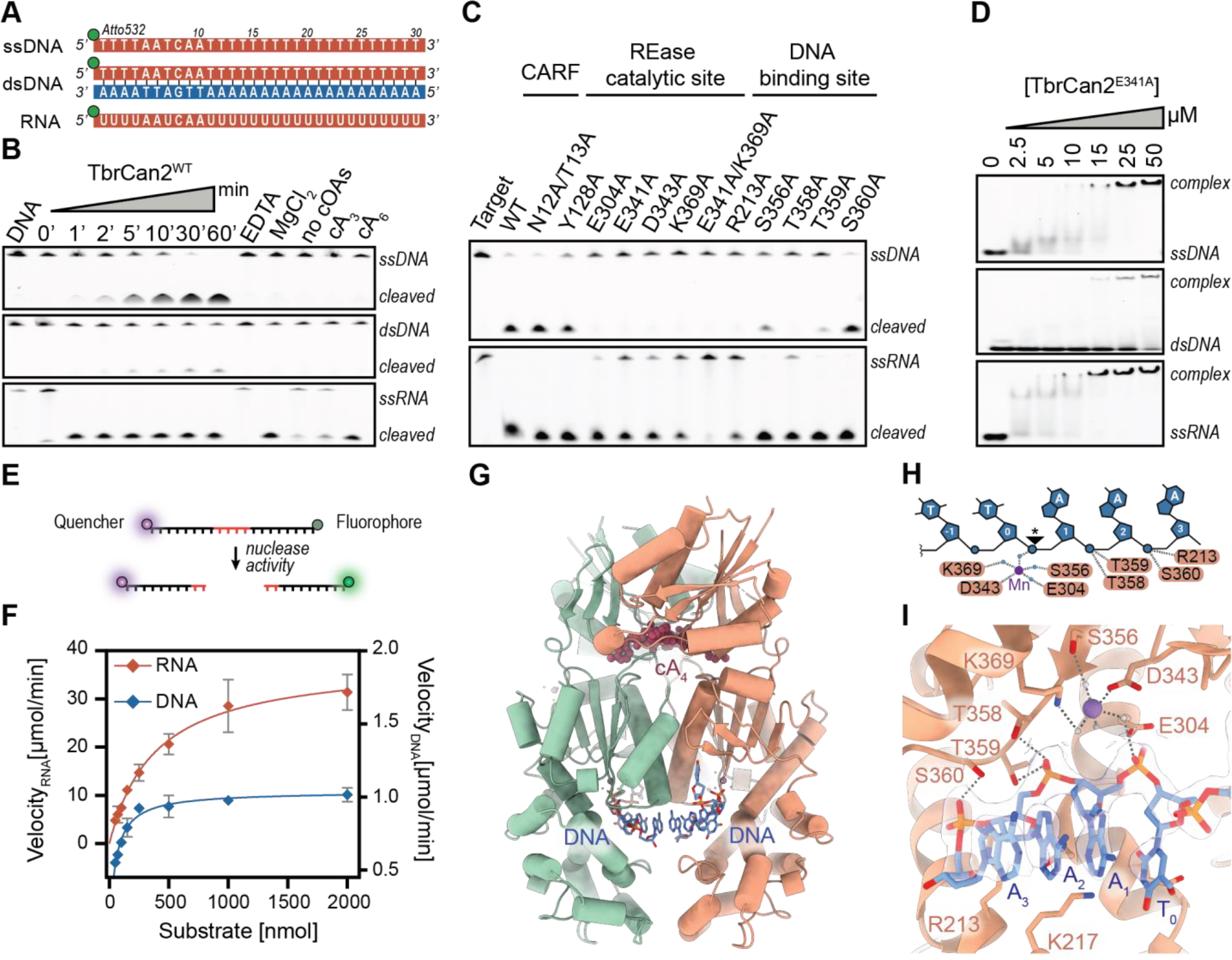
TbrCan2 has ssRNA- and ssDNA-endonuclease activities. (**A**) Schematic representation of nucleic acid substrates used in *in vitro* nuclease assays. (**B**) TbrCan2-catalyzed cleavage of 5’-fluorophore-labeled ssDNA, RNA and dsDNA in the presence and absence of cOAs and MnCl_2_ under single-turnover conditions. (**C**) DNase and RNase activities of structure based mutant TbrCan2 proteins. (**D**) Electrophoretic mobility shift assay of TbrCan2^E341A^ binding to DNA and RNA substrates. (**E**) Schematic of the fluorophore-quencher reporter assay to determine enzyme kinetics of TbrCan2. (**F**) Multiple-turnover kinetics of TbrCan2 on ssDNA and ssDNA substrates. (**G**) Crystal structure of catalytically inactive E341A mutant of TbrCan2 in complex with cA_4_, Mn^2+^ and a ssDNA substrate (5’-TTTAAA-3’) at a resolution of 2.2 Å. (**H**) Schematic diagram of the DNA substrate and the protein-DNA interactions observed in the crystal structure. The scissile phosphate is denotded with an asterisk. Blue dots represent water molecules. (**I**) Close-up of the REase active site showing a bound substrate. 2m*F*_0_-D*F*_C_ composite omit map in **I** is contoured at 1.0 σ and displayed within a radius of 2 Å around the DNA.

Next, we assessed the binding affinities of TbrCan2 for the oligonucleotide substrates by electrophoretic mobility shift assays (EMSAs), providing further evidence that TbrCan2 primarily functions as a single-strand selective nuclease. Furthermore, we observed that TbrCan2^E341A^ bound to DNA and RNA with apparent dissociation constants (K_D_) of 2.9 μM and 0.24 μM, respectively, suggesting a higher overall binding affinity for RNA (**Figure 2D**). This was corroborated by complementary multiple-turnover kinetics experiments (**Figure 2E**), in which the observed rate constant for ssRNA cleavage (k_cat_ = 7.6 min^-1^) was approximately two orders of magnitude higher than that for ssDNA cleavage (k_cat_ = 0.1 min^-1^), indicating a strong selectivity for ssRNA substrates by TbrCan2 (**Figure 2F**).

### Substrate binding mechanism of TbrCan2

We sought to elucidate the substrate binding mechanism of TbrCan2 by determining the structure of TbrCan2 in complex with a ssDNA substrate. To this end, we initially solved a 2.5 Å structure of the catalytically inactive E341A mutant (TbrCan2^E341A^) in complex with cA_4_ and a 5’-TTTAAA-3’ ssDNA (**Figure 2G-I**). The TbrCan2-cA_4_ complex symmetrically binds two DNA molecules within the REase active sites which face a central substrate binding channel (**Figure 2G**). The bases of nucleotides dT0 to dA3 of both strands form a continuous π-π stack, with the 5’-terminal nucleotides facing the binding channel exit. The scissile phosphate group located between nucleotides dT0 and dA1 is positioned near the Mn^2+^ ion, hydrogen bonding to Glu304 and a metal-coordinating water molecule (**Figure 2H, I**). DNA binding is mediated by hydrogen bonding interactions between the phosphate groups of nucleotides dA2 and dA3, and side chain hydroxyl groups of Thr358, Thr359 and Ser360 from the conserved SxTTS motif (**Figure 2H, I**).

Substrate binding is further stabilized by salt bridges between the scissile phosphate and Lys301, and between the dA3 phosphate and Arg213. In agreement with these observations, TbrCan2 nuclease activity was disrupted by mutations R213A, T358A and T359A and significantly reduced for S356A and S360A mutants, highlighting the importance of these residues for substrate nucleic acid binding (**Figure 2C and Supplementary Figure S5**).

The TbrCan2-ssDNA complex structure revealed that the two REase active sites in the TbrCan2 homodimer can engage two DNA strands simultaneously (**Figure 2G**), yet TbrCan2 preferentially cleaves single-stranded nucleic acids and exhibits only moderate nickase activity in the presence of dsDNA substrates (**Figure 1F**). To gain insight into the single-stranded target selectivity, we modeled a B-form DNA duplex into the TbrCan2 active site using the ssDNA-bound TbrCan2 structure as a reference. Positioning the substrate strand of the dsDNA duplex in the REase active site based on superposition with the 5’-TTTAAA-3’ ssDNA resulted in steric clashes of the non-substrate strand with the protein backbone of both TbrCan2 protomers (**Supplementary Figure S6**). This explains the lack of dsDNA endonuclease activity in TbrCan2 and implies that dsDNA binding and nicking may require conformational changes in TbrCan2 or structural distortions in the bound dsDNA. Attempts to elucidate the disparities in double- and single-strand selective DNase and RNase activities by co-crystallizing TbrCan2 with dsDNA or RNA substrates remained unsuccessful, potentially due to the low binding affinities for dsDNA substrates and the residual RNase activity observed for TbrCan2E341A (**Supplementary Figure S5**).

### Base selectivity for CA dinucleotides

The nuclease domain of Can2 nucleases shares similarities with restriction endonucleases, which exhibit strong sequence specificity. Previous studies have reported that Can2 nucleases show a certain bias for nucleotide sequences (26, 28). However, the structure of the TTTAAA ssDNA-bound complex did not reveal specific contacts between the protein and the bases of the DNA substrate. To test for sequence selectivity of TbrCan2, we performed nuclease activity assays in the presence of 15-mer oligo-A, - C, and -U/T nucleotides. TbrCan2 efficiently bound and degraded rA_15_ RNA, while it showed reduced turnover and binding of rC_15_ RNA (**Figure 3A-B and Supplementary Figure S7a-b**). In contrast, rU_15_ RNA, dT_15_ DNA or G-containing DNA (T_7_-G_3_-T_20_) substrates were not cleaved in *in vitro* cleavage assays, nor were they stably bound by TbrCan2 in EMSA binding assays. Altogether, these results suggest a base preference of TbrCan2 for A and C nucleotides (**Figure 3A, B**).

**Figure 3.**
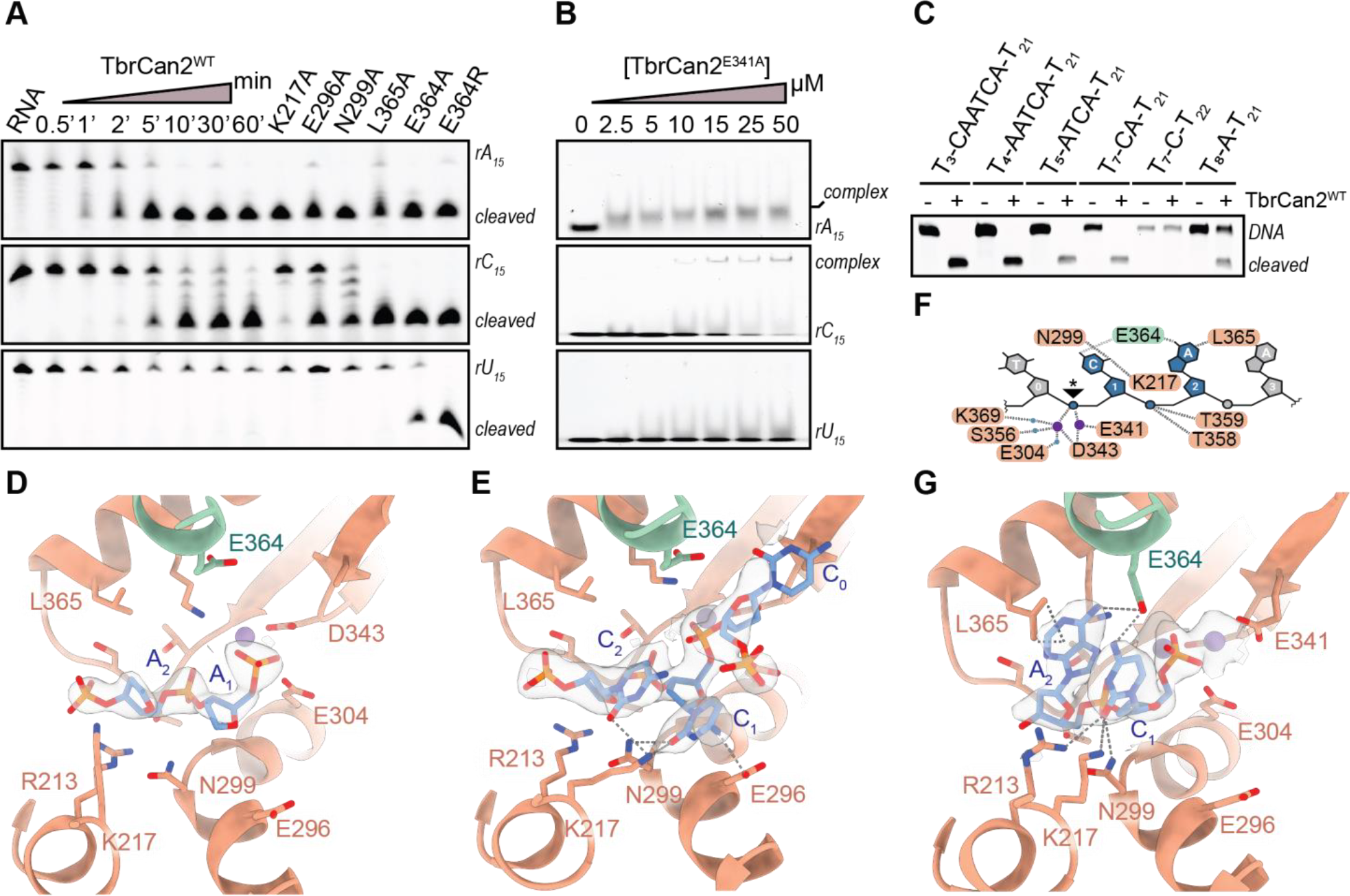
TbrCan2 displays preference for CA dinucleotides. (**A**) Nuclease activity assay of WT and mutant TbrCan2 enzymes with oligo-A, -C and -U RNA. (**B**) EMSA binding assays of TbrCan2^E341A^ with RNA substrates. (**C**) Cleavage of different DNA of varying sequence by TbrCan2^WT^. (**D**) Crystal structure of dA_6_-bound TbrCan2^E341A^-cA_4_ complex, determined at a resolution of 2.0 Å. (**E**) TbrCan2^E341A^-cA_4_ in complex with dC_6_ DNA, determined at a resolution of 2.2 Å. (**F**) Schematic depicting interactions responsible for substrate binding by wild-type TbrCan2. (**G**) Crystal structure (2.2 Å) of the post-catalytic TbrCan2-cA_4_ complex bound to a 5’-pCpAp-3’ dinucleotide cleavage product. The 2m*F*_0_-D*F*_C_ composite omit maps in **D, E** and **G** are shown contoured at 1.0 σ within a radius of 2 Å around the DNA.

Next, we tested the nuclease activity of TbrCan2 on ssDNA substrates designed to contain various sequence motifs (**Figure 3C and Supplementary Figure S7d, e**). We observed efficient degradation of sequences containing CA di-nucleotide motifs (**Figure 3C**), whereas cleavage of DNAs containing only A or C nucleotides was significantly slower (**Supplementary Figure S7d, e**). TbrCan2 did not cleave AC, GC or CG motifs, suggesting that efficient cleavage requires the recognition of a CA di-nucleotide motif (**Supplementary Figure S7d, e**). To identify residues involved in the base selectivity, we solved the crystal structures of TbrCan2^E341A^-cA_4_ complex bound to dA_6_ and dC_6_ DNA oligonucleotides at a resolution of 2.0 Å and 2.3 Å, respectively (**Figure 3D, E**). In the dA_6_-bound structure, only the DNA backbone is resolved, with no observable density for the adenine bases, suggesting that the nucleobases do not make specific contacts within the active site (**Figure 3D**). In contrast, four consecutive nucleotides are resolved in the dC_6_-bound structure, revealing a distortion of the DNA backbone around dC0, where the bases are stabilized by hydrogen bonding interactions between the dC1 and dC2 bases and Lys217, Glu296, and Asn299 from the INS domain (**Figure 3E**). Alanine substitutions of these residues substantially reduced rC_15_ cleavage *in vitro*, supporting their involvement in cytosine base interactions, while rA_15_ degradation was only partly reduced (**Figure 3A**). Moreover, the dC_6_-bound structure revealed that the acidic side chain of Glu364 from the opposite protomer protrudes into the active site in proximity to the 4-amino group of the dC nucleobases (**Figure 3F, G and Supplementary Figure S7f**). Intriguingly, the E364A mutant (TbrCan2^E364A^) retained rA_15_ and rC_15_ cleavage activities and in addition degraded rU_15_ and T_7_-G_3_-T_20_ substrates (**Figure 3A and Supplementary Figure S7a**). The E364R mutation further increased the rU_15_ and T_7_-G_3_-T_20_ cleavage rates. The crystal structure of TbrCan2^E364R^ bound to dT_6_ DNA reveals that the sidechain of Arg364 is within hydrogen bonding distance of the carbonyl O4 atoms of the dT2 base (**Supplementary Figure S7g**). Together, these observations point to Glu364 as a major determinant of the nucleobase preference of TbrCan2.

The structures of the substrate-bound complexes of the catalytically inactive mutant TbrCan2^E341A^ reveal only one Mn^2+^ bound in the REase active site. In consequence, the observed mode of substrate binding might be perturbed due to the absence of the second divalent cation. To gain insights into nucleic acid binding in the context of the wild-type enzyme, we co-crystallized wild-type TbrCan2 (TbrCan2^WT^) in the presence of cA_4_, Mn^2+^ ions and a CA-containing oligonucleotide (5’-AATCAATCA-3’ ssDNA). The resulting structure, determined at a resolution of 2.2 Å, captures the TbrCan2^WT^-cA_4_ complex in a post-catalytic state, bound to dCdA dinucleotide 3’-terminal cleavage products (**Figure 3F**). The nucleobases of the cleavage product are recognized through hydrogen bonding interactions between the 6-amino group of dC0 and the 4-amino group of dA1 with Glu364 in the opposite TbrCan2 protomer. These observations confirm the role of this amino acid residue in the base selectivity of TbrCan2 (**Figure 3F**). Interestingly, the orientation of the dC_6_ substrate DNA within the DNA binding channel is distinct from that of the dCdA di-nucleotide cleavage product (**Supplementary Figure S7c**). It remains uncertain whether these differences reflect distinct pre- and post-catalytic binding modes, differential binding of CA-containing nucleotides, as opposed to oligo-C DNA, or are a consequence of the E341A mutation and absent Mn^2+^ ion in the dC_6_-bound substrate complex. Taken together, the biochemical and structural data demonstrate that TbrCan2 has sequence preference for CA dinucleotides located immediately downstream of the scissile site, which can be attributed to direct interactions with the CA nucleobases.

### Nuclease activation by cA_4_-induced rearrangement of the catalytic site

Previous studies of CARD1 have revealed that cA_4_ binding induces global conformational changes that bring the REase domains into closer proximity, which was postulated to result in catalytic activation (28). However, the precise mechanism of cA_4_-induced activation of Can2/CARD1 enzymes remains elusive. To address this question, we additionally determined a crystal structure of TbrCan2^WT^ in the apo state, at a resolution of 2.0 Å, to allow for comparisons with the structures of TbrCan2^WT^-cA_4_ and the product-bound TbrCan2^WT^-cA_4_ complex (**Figure 4A-C**). Comparison of the apo and cA_4_-activated structures reveals pronounced conformational changes whereby the two REase domains undergo a shearing motion (**Figure 4A, B and Supplementary Figure S8a**), similar to the structural changes observed in TresuCARD1 upon cA_4_ binding (28). In addition to this global structural rearrangement, we observe marked local changes of beta-strand 10 (β10), which contains catalytic residues Glu341 and Asp343 within the ExD-SxTTS-K REase motif. As β10 strand moves within the active site, Glu341 is shifted by approximately 1.5 Å towards the metal binding cavity (**Figure 4B and Supplementary Figure 8b**). This restructuring of the active site enables the binding of a second catalytic Mn^2+^ ion by bringing Glu341 within inner-coordination distance of the Mn^2+^_b_ ion, while Asp343 is now positioned to simultaneously coordinate the Mn^2+^_a_ and Mn^2+^_b_ cations (**Figure 4C, F**). Catalytic activation is allosterically coupled to cA_4_ binding through a cascade of interactions. A loop comprising residues 382-385 (**Figure 1H**) that connects α-helix 15 (α15) and β12 of the REase domain protrudes into the cOA binding cavity formed by the two CARF domains (**Figure 4E**). The binding of cA_4_ is sensed by interactions between the A_1_/A_3_ base and Asp383 and Glu384, thereby pushing onto the loop, which repositions α15 within the REase domain. This movement is transmitted onto β10 through interactions between Ile380 (α15) and Phe340 (β10), thereby positioning Glu341 and Asp343 for Mn^2+^ binding. The active conformation of TbrCan2 is further stabilized by placing Arg376 (α15) within salt-bridging distance of Asp404 (α15) of the opposite protomer as the two REase domains move into proximity. Mutations of residues that are involved in this cascade abolish Can2 activation and impair nuclease activity (**Figure 4G**), thus supporting the allosteric activation model.

**Figure 4.**
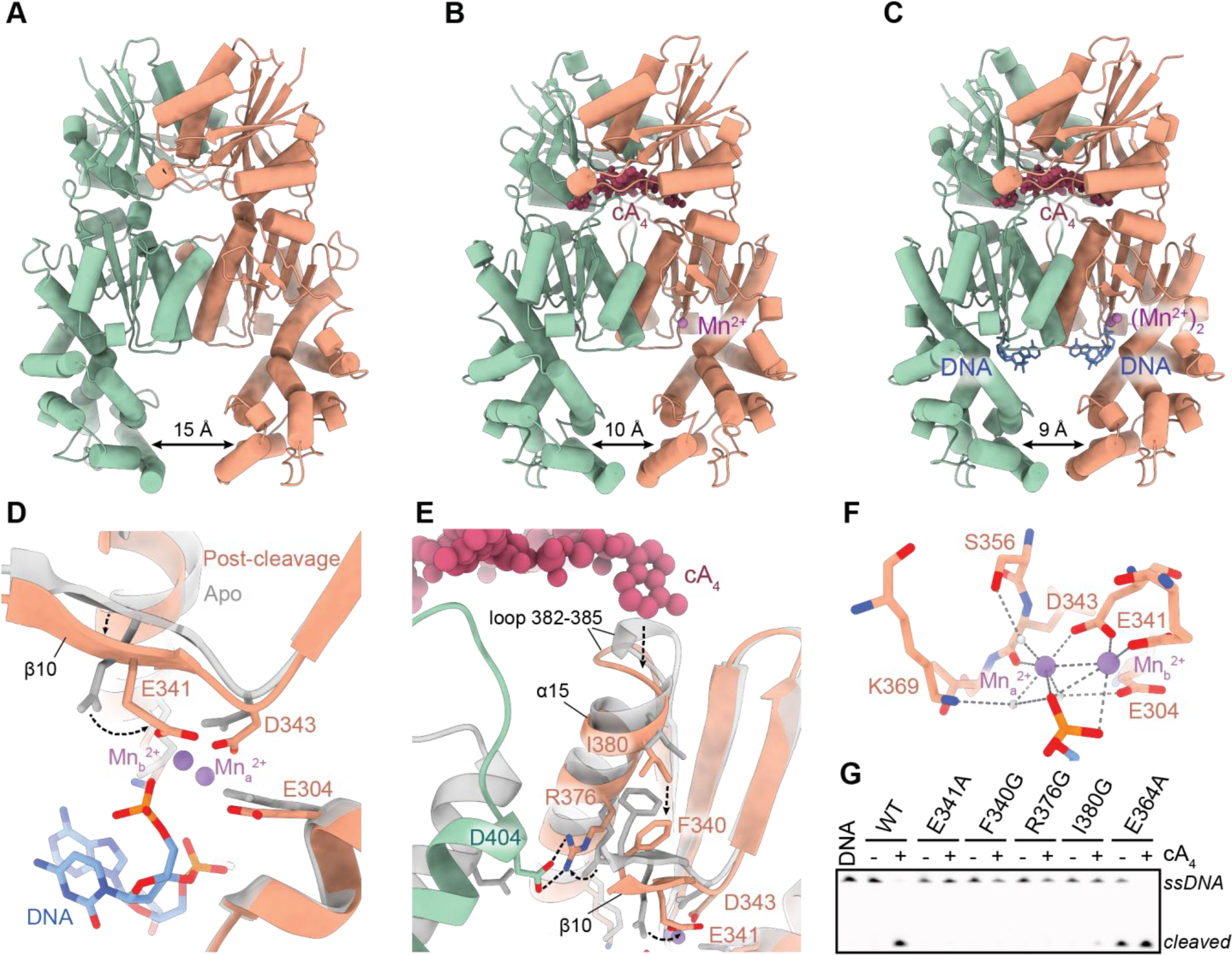
cA_4_-induced structural rearrangement of divalent cation binding sites underpins catalytic activation of TbrCan2. (**A-C**) Crystal structures of (**A**) apo TbrCan2^WT^, (**B**) cA_4_-bound TbrCan2^WT^ and (**C**) TbrCan2^WT^ in the post-catalytic state with a 5’-pCpAp-3’ DNA fragment bound to the active site. (**D-E**) Close-up views of the REase active site depicting local conformation transitions induced by cA_4_ binding. gray: apo; colored: post-catalytic. (**F**) Coordination of two catalytic metal ions within the REase active site. (**G**) ssDNA cleavage activity of TbrCan2 active site mutants of residues involved in the cA_4_-mediated activation cascade.

## DISCUSSION

The cyclic oligoadenylate signaling mechanism of type III CRISPR-Cas systems is used to activate a broad spectrum of ancillary proteins, including CARF domain-containing effector nucleases (13, 43–45). As the nuclease activity of these enzymes leads to growth arrest and cell death upon persistent activation of the type III CRISPR-Cas system (28, 46), this necessitates stringent control of the cOA-dependent activation mechanism. Although previous structural studies have captured CARF-domain nucleases in the inactive and cOA-bound active conformational states, the details of the activation mechanism have remained elusive due to the lack of structural information on substrate binding and cleavage.

In this study, we provide critical insights into the nucleic acid binding and cA_4_-dependent catalytic activation mechanism of the type III-D ancillary nuclease TbrCan2. We show that TbrCan2 is a cA_4_-dependent, dual-specificity PD-(D/E)xK nuclease, capable of cleaving single-stranded DNA and RNA substrates through a two-metal-ion catalytic mechanism. Our structural analysis points to a mechanism in which cA_4_ binding induces restructuring of the REase active site. This rearrangement is brought about by a cascade of interactions that link the cA_4_ binding site with a key structural element (β-strand 10) in the REase domain that houses the metal-coordinating catalytic residues (**Figure 5A, B**), ultimately resulting in proper positioning of the divalent metal ions for substrate binding and catalysis. Although the specific amino acid residues involved in the allosteric relay are conserved among orthologs from clade 1, they are notably absent from other Can2 clades (**Figure 5A, B**), suggesting a distinct activation mechanism. Previously determined structures of TresuCARD1 in both apo and cA_4_-bound states reveal that the REase loop connecting α-helix 15 and β-strand 14 is pulled towards the cOA binding cavity, resulting in a marked clockwise rotation of α15 during the transition from the inactive to the active state. This movement positions a conserved lysine residue, which is involved in coordinating a water molecule between the scissile phosphate and one of the two metal ions (**Figure 5C**). The active state conformation of α15 is stabilized through hydrophobic interactions with residues from the opposite protomer and enabled by the rotation and approximation of the two REase domains. These structural comparisons suggest that Can2-like auxiliary nucleases have evolved two distinct activation mechanisms: a “push-and-flip” mechanism, exemplified by TbrCan2 from clade 1 (**Figure 5D**), and a “pull-and-rotate” mechanism, as observed in the clade 5 ortholog TresuCARD1 (**Figure 5E**).

**Figure 5.**
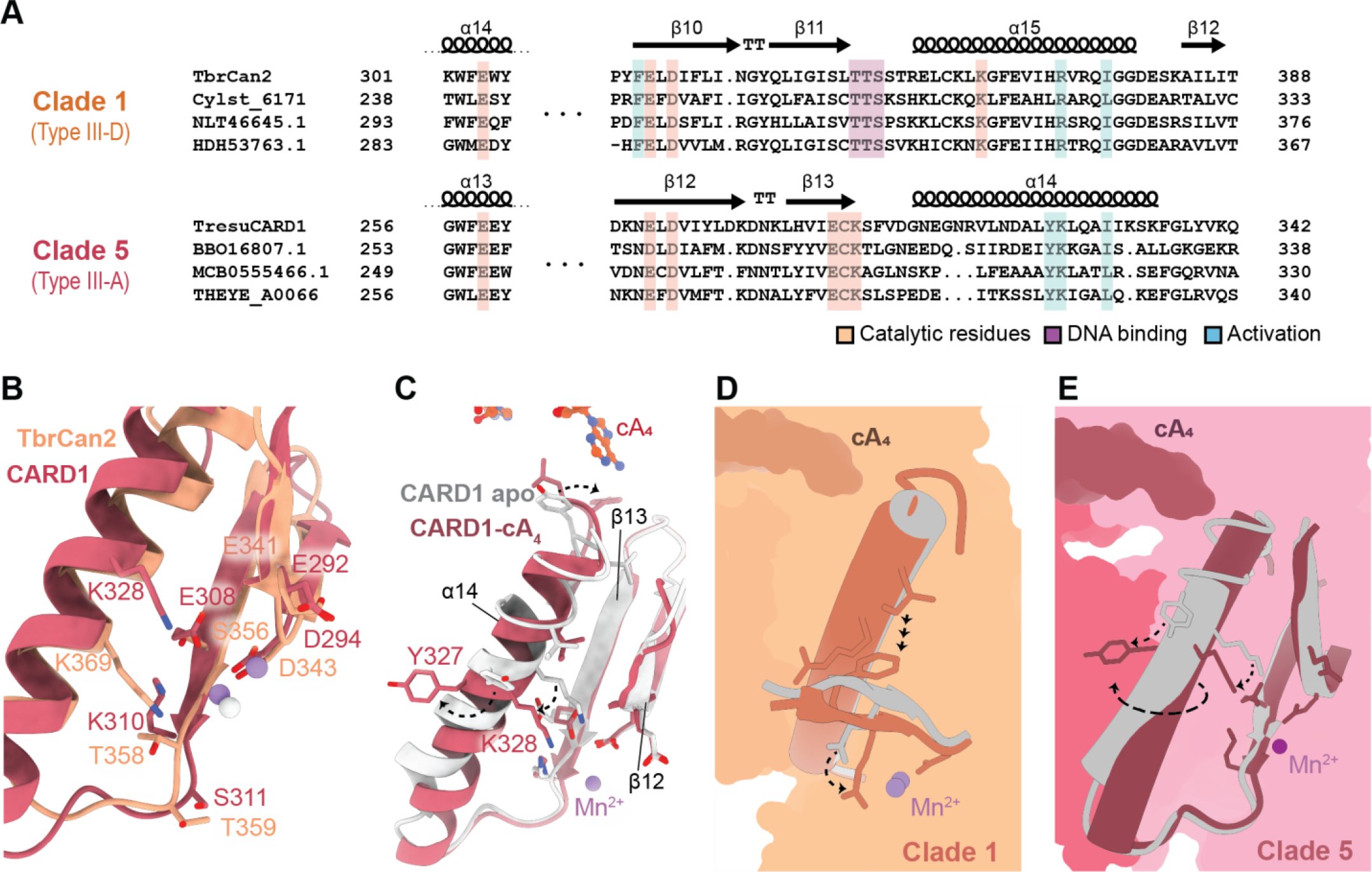
Allosteric activation mechanisms of Can2-family nucleases. (**A**) Sequence alignments of clade 1 and clade 5 Can2 enzymes. Strictly conserved active site residues involved in catalysis (orange), DNA binding (purple) and cA_4_-mediated activation (green) are highlighted. (**B**) Superposition of active site residues in TbrCan2 (orange) and TresuCARD1 (magenta). (**C**) Superposition of apo- and cA_4_-bound TresuCARD1 structures reveals the rotation of α-helix 14 upon cA_4_ binding, potentially activating the REase domain. (**D-E**) Schematic representation of the conformational activation mechanisms of (**D**) clade 1 (TbrCan2) and (**E**) clade 5 Can2 nucleases (TresuCARD1).

A key feature of TbrCan2 is its selectivity for RNA and DNA substrates. The substrates are bound within a channel located between the two REase domains, predominantly lined with charged side chains. The spatial disposition of the two REase active sites enables the concurrent coordination of two DNA or RNA strands. The proper orientation of the substrate is ensured by a conserved SxTTS motif that binds to the 3’ end of the target nucleic acid, positioning the scissile phosphate within the catalytic site. The coordination of the substrate is further secured through interactions with active site residues and metal ions. Intriguingly, the SxTTS motif is absent in Can2 clades other than clade 1, indicating potential mechanistic differences in substrate coordination and likely reflecting differential substrate specificities observed in previously characterized Can2 orthologs (26–28).

The structures of TbrCan2 bound to DNA further rationalize the observed substrate selectivity for CA dinucleotide sequence motifs by hydrogen bonding interactions with Glu364, while T and U bases are discriminated by the absence of productive hydrogen bonding with Glu364 and its negative charge. In agreement with the structural observation, charge reversal at this position confers substrate preference for U- and G-containing substrates. Importantly, numerous clade 1 orthologs feature an arginine residue in the equivalent position, implying a distinct base preference. While the substrate selectivity of TbrCan2 is not as stringent as the sequence specificity displayed by PD-(D/E)xK type II restriction endonucleases, some similarities exist between the two. For example, the restriction enzyme NgoMIV targets a conserved 5’-GCCGGC-3’ hexanucleotide sequence, whose recognition is facilitated by an Arg-Ser-Asp-Arg (RSDR) amino acid motif located near the active site, at a position analogous to α15 in TbrCan2 (**Supplementary Figure S9**) (47). In NgoMIV, the RSDR residue Asp193 connects the amino groups of the C bases of the DNA target strand within the opposing REase active site, a function comparable to Glu364 in TbrCan2 (**Supplementary Figure S9c**), highlighting the silimilarities between substrate recognition mechanisms of restriction enzymes and Can2. However, TbrCan2 lacks the extensive network of interactions that enable specific sequence recognition in NgoMIV, thus resulting in less stringent substrate selectivity. Another property that sets TbrCan2 apart from PD-(D/E)xK domain restriction endonucleases is its selectivity for single-stranded RNA and DNA substrates. While TbrCan2 can bind to double-stranded DNA with low affinity and can catalyze nicking at a low rate, it is incapable of generating double-strand breaks. In type II restriction endonucleases such as EndoMS and NgoMIV, both active sites bind and cleave respective strands simultaneously (**Supplementary Figure S9b**) (47, 48). In contrast, structural modeling of dsDNA binding to TbrCan2 results in steric clashes between dsDNA duplex and the protein backbone, indicating that TbrCan2 is unable to catalyze staggered nicks in a dsDNA substrate due to the specific arrangement of its two catalytic sites and the orientation at which the substrate DNA is coordinated by the SxTTS motif in each REase active site.

We have been unable to determine the structure of TbrCan2 in complex with an RNA substrate, necessitating further investigations into the mechanisms underlying the ribonuclease and deoxyribonuclease activities of CARD1/Can2 enzymes. This is particularly important in light of contradictory reports of the *in vitro* and *in vivo* nuclease activities of TresuCARD1 (28). Although CARD1, like TbrCan2, cleaves RNA targets more rapidly than DNA *in vitro*, its expression in a heterologous host did not affect RNA transcript levels. Although this would suggest that CARD1-mediated growth inhibition might primarily be due to the ssDNAse activity, the relative contribution of the DNAse and RNase activities to the functional mechanism of CARD1/Can2 enzymes is presently unclear. Future work will thus be needed to define the functional landscape of CARD1/Can2 accessory nucleases so as to comprehend their biological functions in their native hosts.

## DATA AVAILABILITY

All data is provided in full in the results section and the Supplementary Data accompanying this paper. Atomic coordinates and structure factors have been deposited in the Protein Data Bank under accession codes 8Q3Y, 8Q3Z, 8Q40, 8Q41, 8Q42, 8Q43 and 8Q44.

## ACKNOWLEDGEMENTS

We would like to thank Beat Blattmann (Protein Crystallization Center, UZH) for the crystallographic support. We also thank Vincent Olieric and Takashi Tomizaki (Swiss Light Source, Paul Scherrer Institute, Villigen, Switzerland) for the assistance with crystallographic data collection. We thank members of the Jinek laboratory for helpful discussions and critical reading of the paper.

## SUPPLEMENTARY DATA

**Supplementary Table S1.**
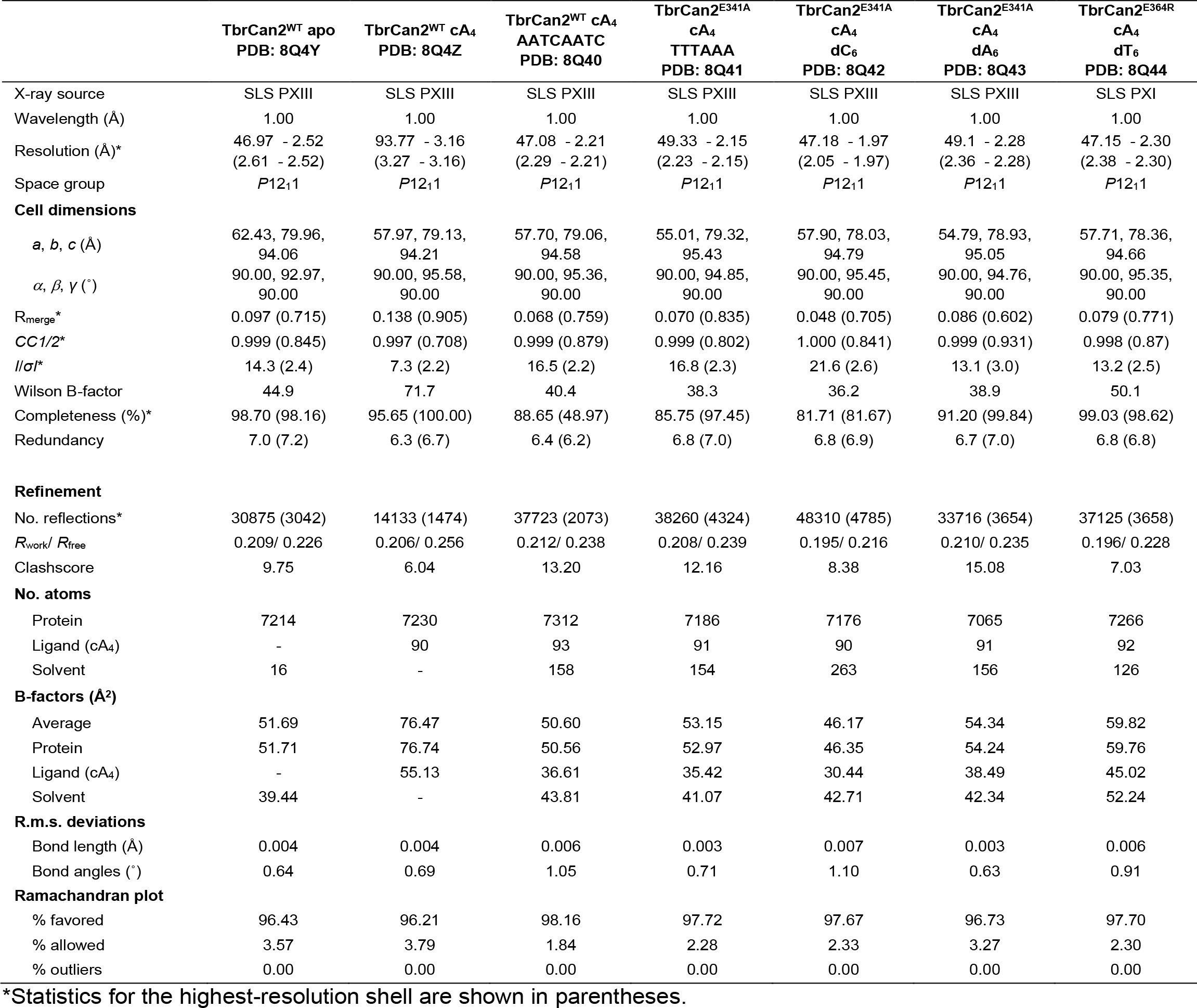
Data collection and refinement statistics.

**Supplementary Figure S1.**
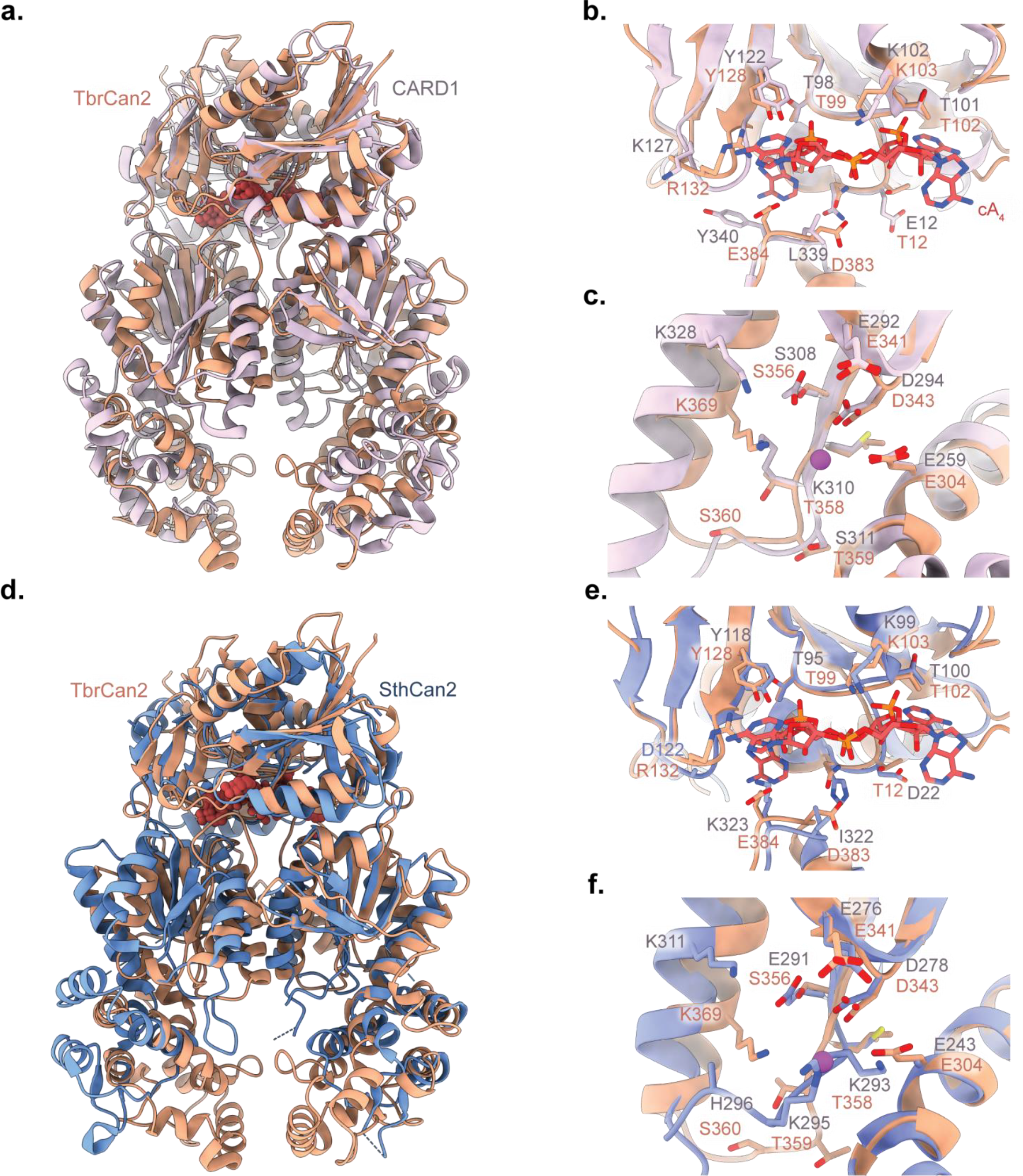
Structural comparison of TbrCan2 and Can2 orthologs. (**a-c**) TbrCan2 aligned with TresuCARD1 in complex with a cA_4_ ligand (PDB: 6wxx). Close-up views of (**b**) the CARF and (**c**) REase active sites in TbrCan2 (orange) and TresuCARD1 (purple). (**d-f**) Structural superposition of TbrCan2 and SthCan2-cA_4_ (PDB: 7bdv). Close-up views of (**e**) the CARF and (**f**) REase active sites in TbrCan2 (orange) and SthCan2 (blue).

**Supplementary Figure S2.**
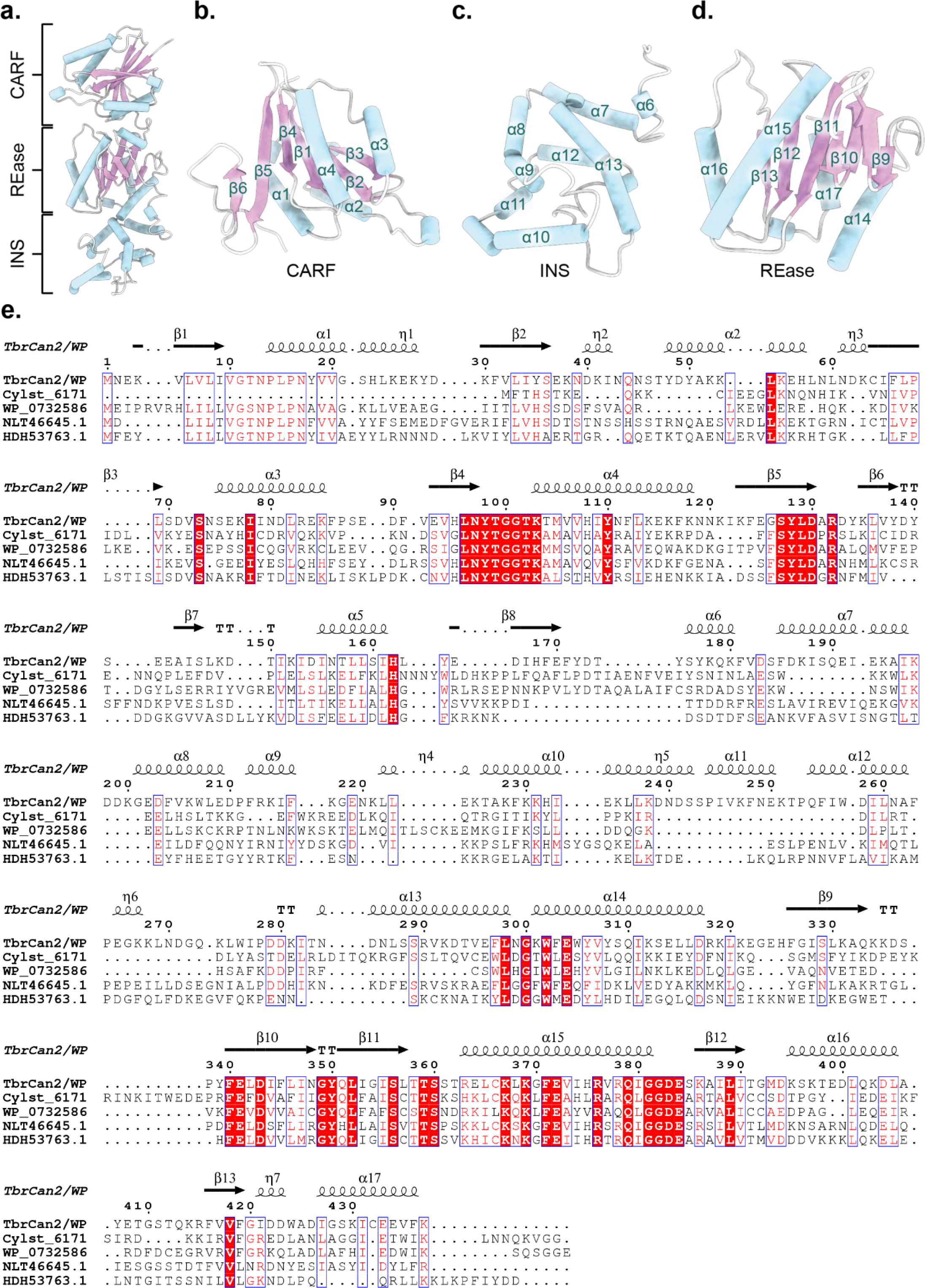
Structure and sequence conservation of clade 1 Can2 orthologs. (**a-d**) Stereo ribbon diagram of TbrCan2. Domains and major secondary structure elements of TbrCan2 are labeled. (**e**) Sequence alignment of TbrCan2 and orthologs from clade 1. Residues of the conserved E-ExD-SxTTS-K motif are located in α14, β10-11 and α15 of the REase domain (Glu304, Glu341, Asp343, Ser356, Thr358, Thr359 and Lys369). Alignment and structure assignment visualized with ESPript (49). Secondary structure elements are indicated above the sequence.

**Supplementary Figure S3.**
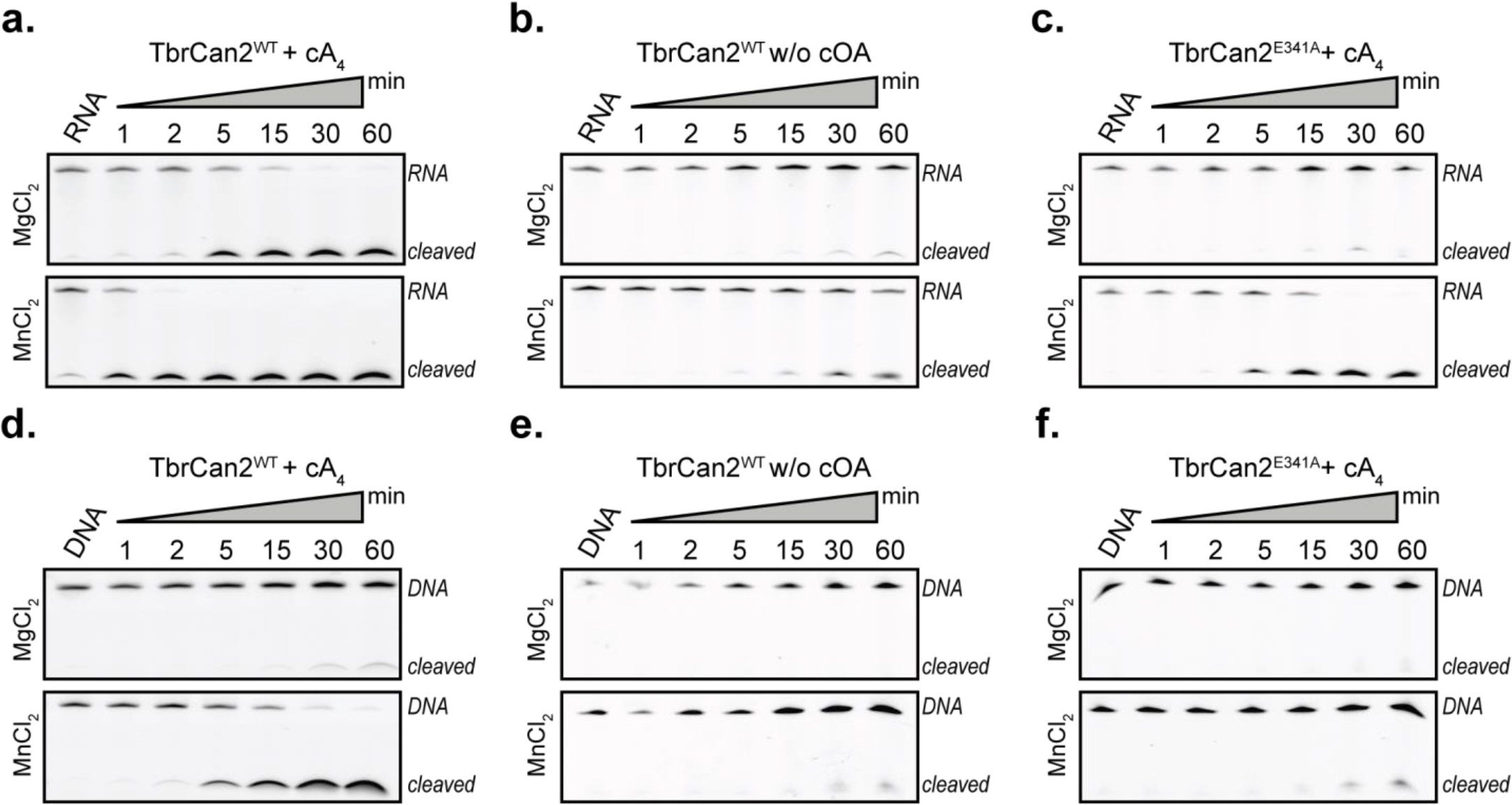
Characterization of TbrCan2 nuclease activity. Cleavage of single-stranded RNA and DNA substrates with WT and E341A TbrCan2 in the presence and absence of cA_4_ and the presence of MgCl_2_ or MnCl_2_.

**Supplementary Figure S4.**
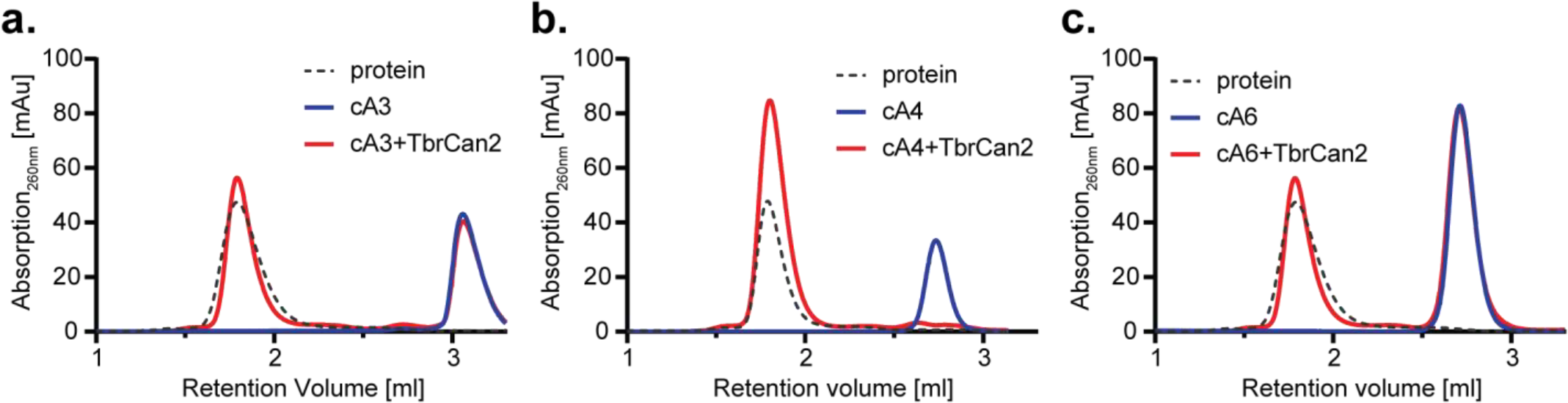
cOA binding assays. Binding of TbrCan2 to (**a**) cA_3_, (**b**) cA_4_ and (**c**) cA_6_ was assayed by analytical size-exclusion chromatography. Black trace: protein; blue: cOA; red: protein + cOAs.

**Supplementary Figure S5.**
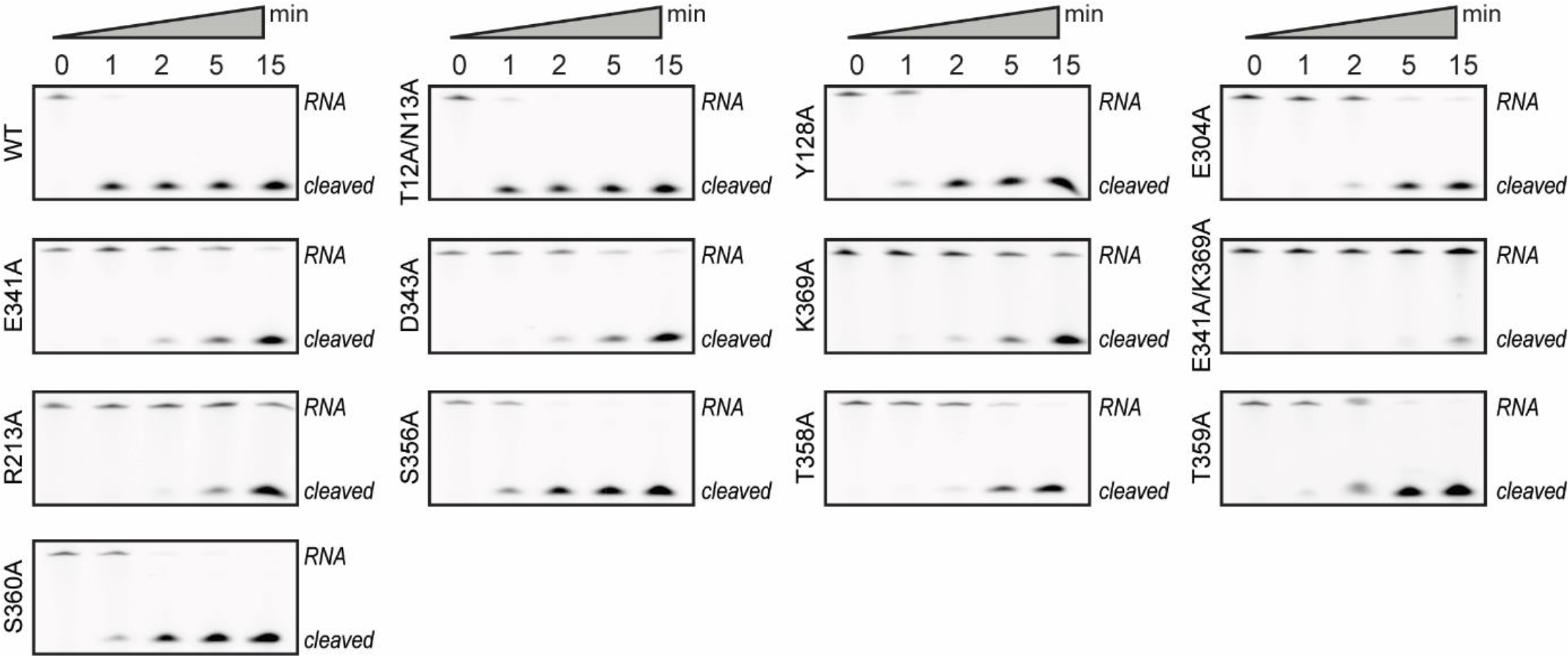
RNase activities of TbrCan2 active site mutants. Nuclease assays of TbrCan2^Mut^ performed with 5’-fluorophore labeled ssRNA substrate under single-turnover conditions.

**Supplementary Figure S6.**
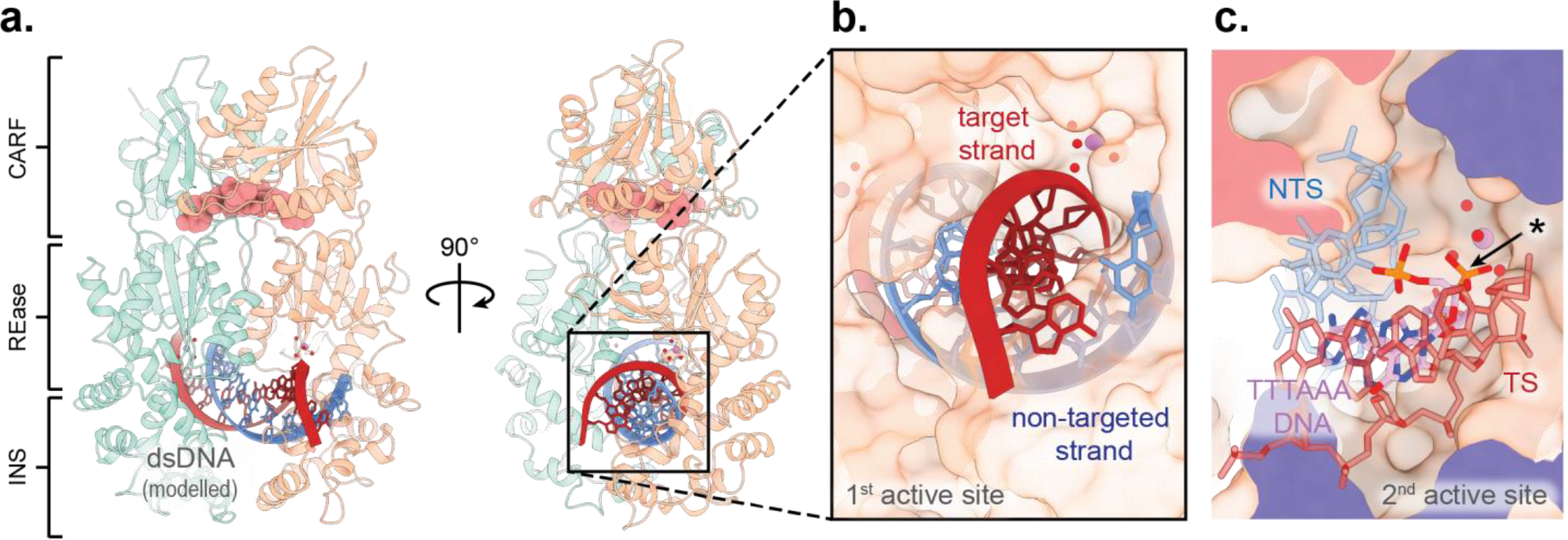
Model of dsDNA binding by TbrCan2. (**a-c**) Structural model showing binding of B-form DNA duplex in the active site of TbrCan2, based on superposition with TTTAAA DNA-bound TbrCan2^E341A^-cA_4_ structure. red: substrate (target) strand (TS); blue: non-target strand (NTS); pink: TTTAAA DNA based on crystal structure. (**b-c**) Close-ups of the (**b**) active site 1, which coordinates the target strand and (**c**) active site 2, which cleaves a second oligonucleotide. *: scissile phosphate.

**Supplementary Figure S7.**
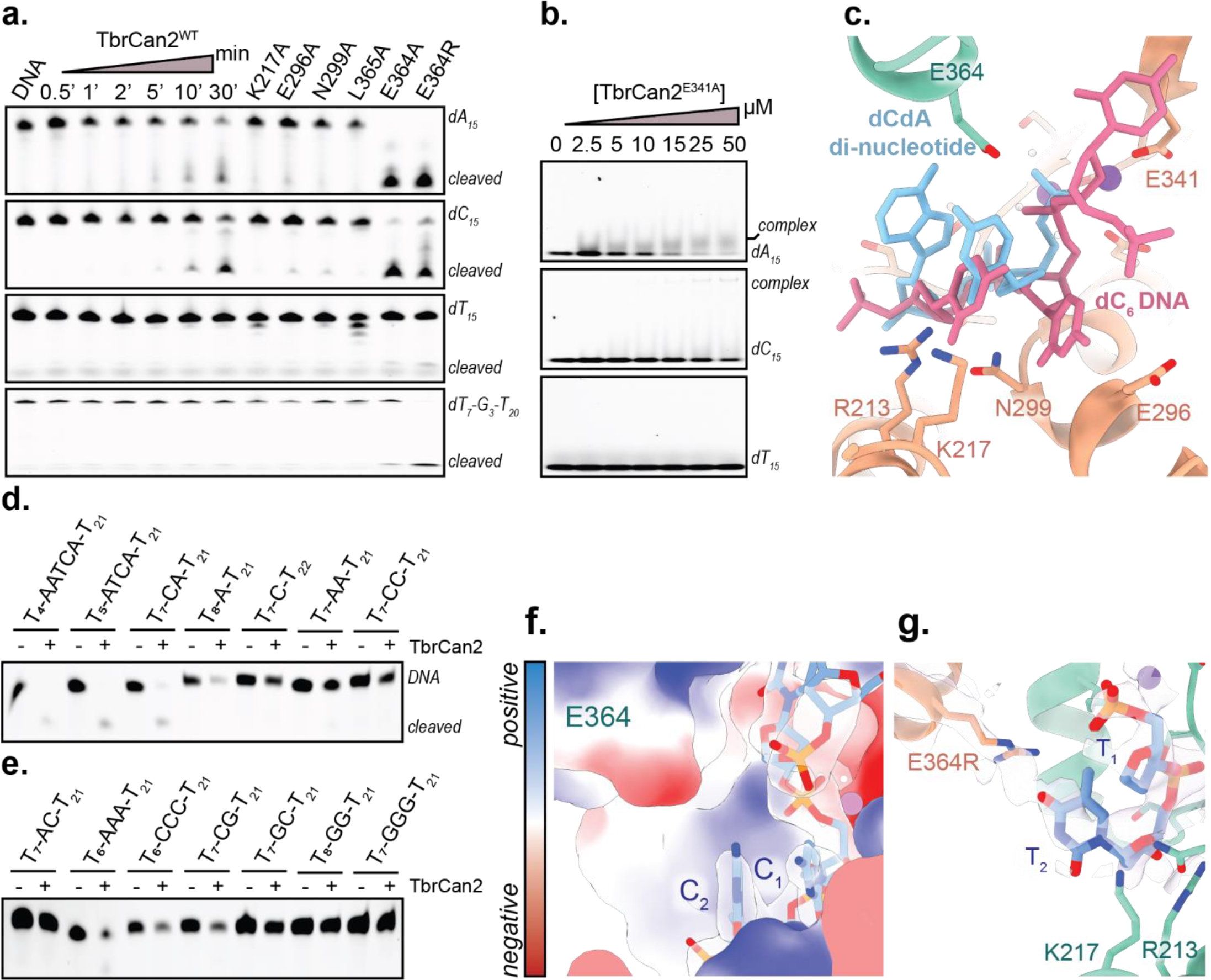
Base-selective binding and cleavage of DNA substrates in TbrCan2. (**a**) Nuclease activity assay of WT and mutant TbrCan2 enzymes with oligo-A, -C and -T ssDNA substrates. (**b**) EMSA binding assays of TbrCan2^E341A^ with DNAs shown in panel **b**. (**c**) Superposition of the dCdA di-nucleotide cleavage product (blue) and the poly-C DNA (pink) within the REase site of post-catalytic TbrCan2^WT^. (**d-e**) Nuclease activity of TbrCan2^WT^ for different DNA substrates with distinct sequence motifs. (**f**) Electrostatic surface charge potential of nucleic acid binding channel in TbrCan2^E341A^, occupied by a dC_6_ oligonucleotide. Coulombic surface coloring range: ± 10 kcal/mol·e; blue: positive, red: negative. (**g**) Crystal structure of T_6_-bound TbrCan2^E364R^-cA_4_ complex at 2.0 Å resolution. Close-up view of the DNA binding site.

**Supplementary Figure S8.**
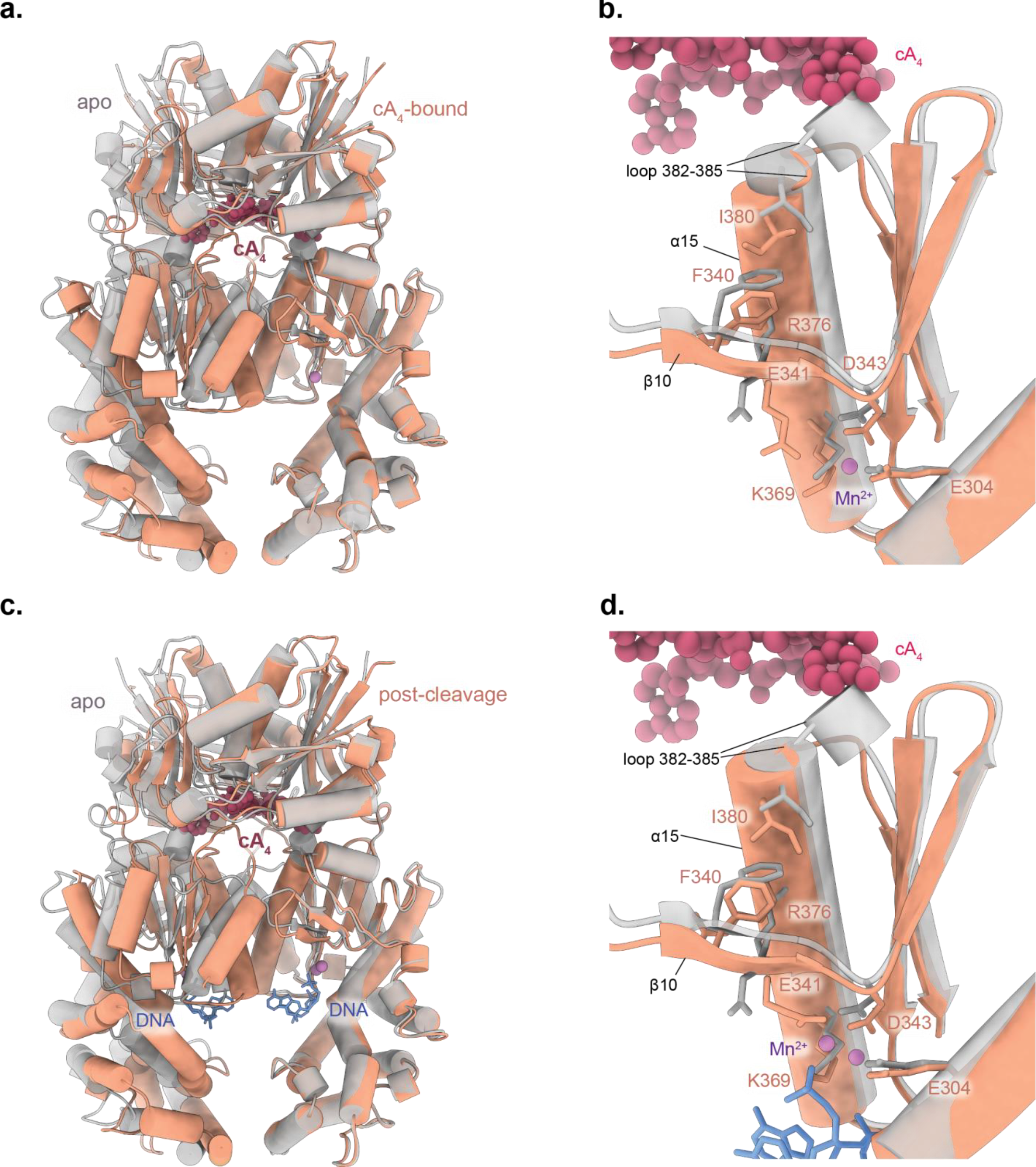
Structural comparisons of Apo and cA_4_-bound TbrCan2. (**a-d**) Structural alignment of apo (gray) TbrCan2 with (**a**-**b**) the cA_4_-bound TbrCan2^WT^ complex (orange) and (**c**-**d**) post-catalytic TbrCan2^WT^ (orange). (**b, d**) Close-ups of the TbrCan2 REase active site.

**Supplementary Figure S9.**
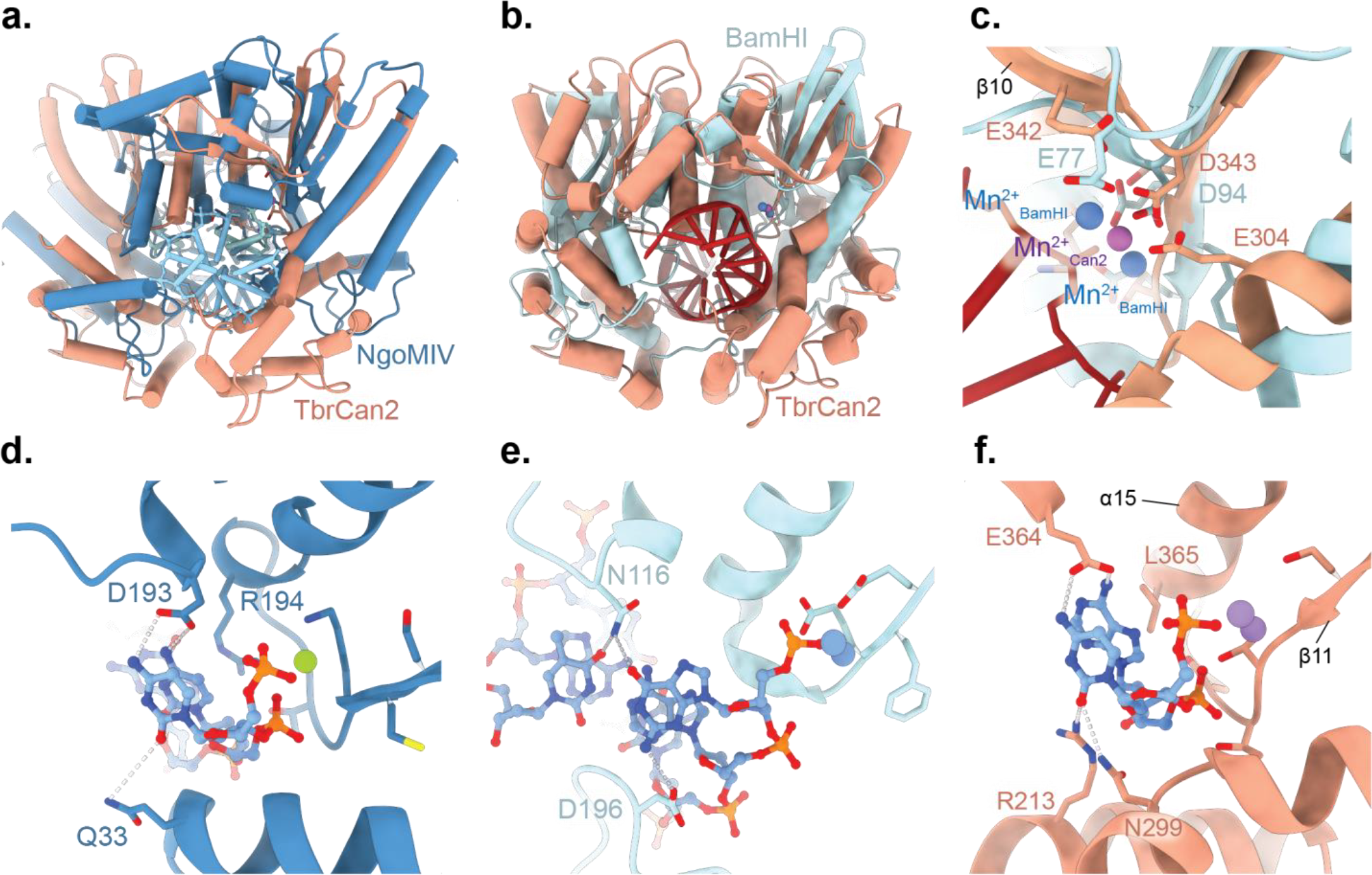
Structural comparisons of substrate recognition in TbrCan2 and type II restriction enzymes. (**a**) Overview of the dsDNA-bound NgoMIV dimer (PDB: 1fiu, blue), aligned with the REase domain of TbrCan2 (orange). (**b**) Overview of the dsDNA-bound BamHI dimer (PDB: 2bam, blue), aligned with the REase domain of TbrCan2 (orange). (**c**) Structural comparison of divalent metal ion coordination in the dsDNA-bound BamHI homodimer (blue) and the TbrCan2^WT^-cA_4_ complex. (**d**-**f**) Close-up views of the DNA binding sites of (**d**) NgoMIV, (**e**) BamHI and (**f**) TbrCan2.

**Supplementary Table S2.** A list of plasmids used in this study.

| Plasmid # | Plasmid name | Source Organism | Resistance | Protein Sequence Source |
| --- | --- | --- | --- | --- |
| pKJ64 | pET16B-6xHis-TEV-TbrCan2(aa1-437) | *Thermoanaerobacter brockii finnii* Ako 1 | AmpR | GenBank: ADV79523.1 |
| pKJ138 | pET16B-6xHis-TEV-TbrCan2D343A(aa1-437) | *Thermoanaerobacter brockii finnii* Ako 1 | AmpR | GenBank: ADV79523.1 |
| pKJ139 | pET16B-6xHis-TEV-TbrCan2E364A(aa1-437) | *Thermoanaerobacter brockii finnii* Ako 1 | AmpR | GenBank: ADV79523.1 |
| pKJ140 | pET16B-6xHis-TEV-TbrCan2E364R(aa1-437) | *Thermoanaerobacter brockii finnii* Ako 1 | AmpR | GenBank: ADV79523.1 |
| pKJ143 | pET16B-6xHis-TEV-TbrCan2T358A(aa1-437) | *Thermoanaerobacter brockii finnii* Ako 1 | AmpR | GenBank: ADV79523.1 |
| pKJ144 | pET16B-6xHis-TEV-TbrCan2T359A(aa1-437) | *Thermoanaerobacter brockii finnii* Ako 1 | AmpR | GenBank: ADV79523.1 |
| pKJ145 | pET16B-6xHis-TEV-TbrCan2E341A(aa1-437) | *Thermoanaerobacter brockii finnii* Ako 1 | AmpR | GenBank: ADV79523.1 |
| pKJ155 | pET16B-6xHis-TEV-TbrCan2N299A(aa1-437) | *Thermoanaerobacter brockii finnii* Ako 1 | AmpR | GenBank: ADV79523.1 |
| pKJ156 | pET16B-6xHis-TEV-TbrCan2Y128A(aa1-437) | *Thermoanaerobacter brockii finnii* Ako 1 | AmpR | GenBank: ADV79523.1 |
| pKJ157 | pET16B-6xHis-TEV-TbrCan2R213A(aa1-437) | *Thermoanaerobacter brockii finnii* Ako 1 | AmpR | GenBank: ADV79523.1 |
| pKJ160 | pET16B-6xHis-TEV-TbrCan2E296A(aa1-437) | *Thermoanaerobacter brockii finnii* Ako 1 | AmpR | GenBank: ADV79523.1 |
| pKJ161 | pET16B-6xHis-TEV-TbrCan2K301A(aa1-437) | *Thermoanaerobacter brockii finnii* Ako 1 | AmpR | GenBank: ADV79523.1 |
| pKJ163 | pET16B-6xHis-TEV-TbrCan2L365A(aa1-437) | *Thermoanaerobacter brockii finnii* Ako 1 | AmpR | GenBank: ADV79523.1 |
| pKJ164 | pET16B-6xHis-TEV-TbrCan2R369A(aa1-437) | *Thermoanaerobacter brockii finnii* Ako 1 | AmpR | GenBank: ADV79523.1 |
| pKJ175 | pET16B-6xHis-TEV-TbrCan2S356A(aa1-437) | *Thermoanaerobacter brockii finnii* Ako 1 | AmpR | GenBank: ADV79523.1 |
| pKJ178 | pET16B-6xHis-TEV-TbrCan2E304A(aa1-437) | *Thermoanaerobacter brockii finnii* Ako 1 | AmpR | GenBank: ADV79523.1 |
| pKJ184 | pET16B-6xHis-TEV-TbrCan2F340G(aa1-437) | *Thermoanaerobacter brockii finnii* Ako 1 | AmpR | GenBank: ADV79523.1 |
| pKJ188 | pET16B-6xHis-TEV-TbrCan2R376G(aa1-437) | *Thermoanaerobacter brockii finnii* Ako 1 | AmpR | GenBank: ADV79523.1 |
| pKJ189 | pET16B-6xHis-TEV-TbrCan2F340G(aa1-437) | *Thermoanaerobacter brockii finnii* Ako 1 | AmpR | GenBank: ADV79523.1 |
| pKJ190 | pET16B-6xHis-TEV-TbrCan2K213A(aa1-437) | *Thermoanaerobacter brockii finnii* Ako 1 | AmpR | GenBank: ADV79523.1 |
| pKJ192 | pET16B-6xHis-TEV-TbrCan2I380G(aa1-437) | *Thermoanaerobacter brockii finnii* Ako 1 | AmpR | GenBank: ADV79523.1 |
| pKJ193 | pET16B-6xHis-TEV-TbrCan2T12A/N13A(aa1-437) | *Thermoanaerobacter brockii finnii* Ako 1 | AmpR | GenBank: ADV79523.1 |
| pKJ196 | pET16B-6xHis-TEV-TbrCan2E341A/K369A(aa1-437) | *Thermoanaerobacter brockii finnii* Ako 1 | AmpR | GenBank: ADV79523.1 |

**Supplementary Table S3.**
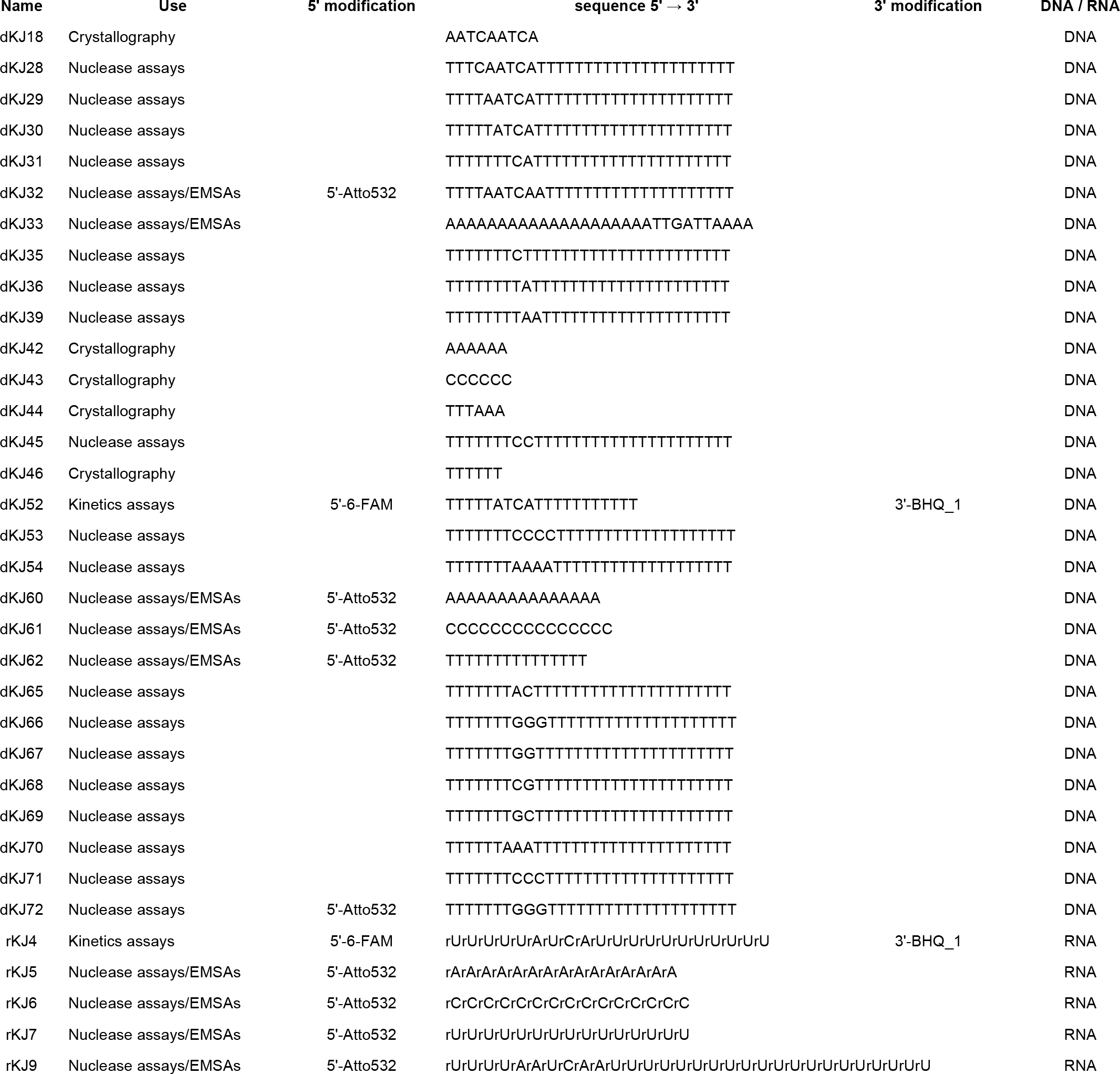
Sequences of oligonucleotide constructs used in this study.

